# Pancreatic cancer extracellular vesicles carry a time-of-day-regulated miRNA cargo that disrupts the skeletal muscle clock and bioenergetics

**DOI:** 10.64898/2026.05.03.722338

**Authors:** Jonathan Church, Ignacio Aiello, Alessandro Ceci, Goeun Jang, Carla V. Finkielstein

**Author notes:** These authors contributed equally (Jonathan Church, Ignacio Aiello). Corresponding author: Carla V. Finkielstein.

## Abstract

Pancreatic ductal adenocarcinoma (PDAC) carries a dismal prognosis, aggravated by cachexia, a systemic wasting syndrome whose molecular mediators remain incompletely defined. Circadian disruption is a further hallmark of PDAC, yet a shared mechanistic basis between these two features has not been established. Here we show that the PDAC secretome, and the small extracellular vesicles (sEVs) it carries, are sufficient to disrupt the circadian clock in independent reporter cell lines and in differentiated myotubes, and to induce myotube atrophy. PANC-1 sEV release and miRNA cargo display pronounced time-of-day variation, with the cargo resolving into two functionally distinct pools: a rhythmically secreted subset, whose release is phase-coordinated and which transmits time-of-day information to the recipient tissue, and a constitutively secreted subset that, although non-rhythmic at the source, is itself capable of perturbing the recipient circadian clock. Individual miRNAs drawn from both pools exert distinct and non-redundant effects on circadian period and myotube diameter. Seahorse extracellular flux analysis further reveals that these miRNAs reprogram mitochondrial respiration and substrate utilization along three divergent trajectories, energetic, high-metabolic, and glycolytic, rather than along a single bioenergetic axis. Intersecting the tumor sEV secretome with serum sEV miRNAs from a pancreatic cancer patient cohort and with the miRNA-Seq landscape of 495 PDAC tumors defines a stable, broadly tumor-abundant, patient-detectable miRNA signature that collectively regulates circadian, proteostatic, and cachexia-relevant gene networks. Across these orthogonal datasets, hsa-miR-27b-3p emerges as a node within the rhythmically secreted pool: consistently detected in patient serum, ranked among the top 50 most abundant miRNAs in >90% of these tumors, and individually sufficient to shorten the circadian period, drive myotube atrophy comparable to dexamethasone, and impose an energetic mitochondrial phenotype. Together, these findings identify PDAC sEV miRNAs as temporally organized mediators coupling circadian disruption, muscle bioenergetics, and cachexia-relevant muscle reprogramming.

## Introduction

Pancreatic ductal adenocarcinoma (PDAC) remains one of the most lethal malignancies, with a five-year relative survival rate of only 13% and an incidence rising at approximately 0.9% per year (Siegel, Kratzer, Wagle, Sung, & Jemal, 2026). Surgical resection, the only potentially curative option, is available to fewer than 20% of patients at diagnosis (Chrystoja et al., 2013), and the available chemotherapeutic regimens, gemcitabine, nab-paclitaxel, FOLFIRINOX, or their combinations, offer only modest survival gains against a backdrop of rapidly emerging chemoresistance (Adamska et al., 2018; Gnanamony & Gondi, 2017). This clinical profile is aggravated by cachexia, a multisystem wasting syndrome that defines advanced disease (Baracos, Martin, Korc, Guttridge, & Fearon, 2018). Cachexia is characterized by involuntary weight loss, systemic inflammation, and skeletal muscle atrophy, and arises from a coordinated imbalance in protein turnover and energy expenditure, driven by the upregulation of catabolic programs (ubiquitin– proteasome, autophagy–lysosomal, calcium signaling), the suppression of anabolic signaling (PI3K/AKT/mTOR), and mitochondrial dysfunction (A. Martin, Gallot, & Freyssenet, 2023; Rausch, Sala, Penna, Porporato, & Ghigo, 2021; Setiawan et al., 2023). Cachexia is not a comorbidity but a cause of death in a substantial fraction of PDAC patients (Baracos et al., 2018), and the mechanisms by which pancreatic tumors reprogram skeletal muscle at a distance remain incompletely defined. Inflammatory mediators have historically dominated the cachexia field (VanderVeen, Fix, & Carson, 2017). Preclinical models implicate TNF-α, IL-6, IL-1β, IL-8, and CRP, yet cytokine levels in patients correlate inconsistently with the clinical features of wasting (Onesti & Guttridge, 2014; Webster, Kempen, Hardy, & Langen, 2020), and trials of anti-TNF-α or anti-IL-6 antibodies have yielded mixed outcomes (Paval et al., 2022). These observations argue that non-cytokine mediators must contribute to the pathogenesis of cachexia (Webster et al., 2020; L. Zhang & Bonomi, 2024).

Small extracellular vesicles (sEVs), lipid-bilayer nanoparticles of endosomal origin released upon multivesicular body fusion with the plasma membrane, have emerged as compelling candidates (Kalluri & LeBleu, 2020). sEVs carry selectively sorted cargo including proteins, lipids, and small non-coding RNAs, and traffic stably through the circulation to reach distal tissues (Kalluri & LeBleu, 2020; O’Brien, Breyne, Ughetto, Laurent, & Breakefield, 2020). Cancer-derived EVs from Lewis lung carcinoma (LLC) and C26 colorectal cancer models promote skeletal muscle catabolism, adipose tissue browning, and lipolysis (Hu et al., 2021; Marzan & Chitti, 2023). The PDAC setting, by contrast, remains comparatively underexplored. While recent work has begun to implicate PDAC-derived sEVs and their miRNA cargo in muscle wasting (Xu et al., 2025; Yang et al., 2019), a systematic characterization of the tumor sEV cargo, of its temporal dynamics, and of its causal contribution to muscle bioenergetic remodeling in cachexia is still lacking (Marzan & Chitti, 2023; Y. Wang & Ding, 2024).

A second, largely separate line of inquiry places muscle bioenergetics at the center of cachectic wasting. Cachectic myofibers exhibit reduced oxidative capacity, impaired mitochondrial biogenesis, and a shift toward glycolytic substrate utilization, changes that precede, and likely precipitate, overt proteolysis (Julienne et al., 2012; Mannelli, Gamberi, Magherini, & Fiaschi, 2021; VanderVeen et al., 2017). These bioenergetic alterations position mitochondrial respiration and substrate flux as functional readouts of the cachectic program rather than as downstream consequences of it.

A third, and until recently unconnected, dimension concerns the circadian clock. Circadian disruption is a well-established feature of PDAC (Relles et al., 2013; Schwartz et al., 2023). Comparative analysis of PDAC and matched normal pancreatic tissue reveals a marked loss of rhythmic coordination among core clock genes, with *BMAL1, PER1, PER3*, and *NR1D1* most severely affected (Schwartz et al., 2023). Low BMAL1 expression correlates with aggressive tumor behavior and reduced overall survival (Li et al., 2016), and *Bmal1* knockout in KPC pancreatic tumor models accelerates growth while enriching proliferative and adhesion-related pathways (Schwartz et al., 2023). *BMAL1* knockdown in BxPC-3 cells likewise enriches growth pathways and suppresses apoptosis (Jiang et al., 2016; Jiang et al., 2018). These findings establish a cell-intrinsic clock defect in PDAC; whether pancreatic tumors are also capable of dismantling the clock in other cell types or tissues, and through which mediators, has not been directly addressed. Importantly, adjacent non-tumor pancreatic tissue retains a functional clock (Huisman et al., 2015), indicating that if tumor-derived signals propagate beyond the primary lesion, they must do so at a distance. Consistent with this possibility, colorectal and lung adenocarcinoma models induce phase alterations in the liver and kidney (Huisman et al., 2015; Masri et al., 2016), even in the absence of metastasis, with IL-6 proposed as a candidate systemic mediator but unlikely to account, on its own, for the full circadian phenotype (Masri et al., 2016).

The intersection of these three threads, sEV-mediated systemic communication, muscle bioenergetic collapse, and tumor-induced circadian reprogramming of peripheral tissues, suggests that cachexia and clock disruption in PDAC may not be independent features of advanced disease but coupled outputs of a shared, sEV-encoded mechanism. miRNAs are among the most abundant and stable components of sEV cargo and offer a parsimonious molecular substrate for this coupling (Rohm, Cunha, & Olefsky, 2025). Computational and experimental work across mouse, zebrafish, and human systems identifies multiple miRNAs that target *Per1, Per2*, and *Per3, BMAL1, CLOCK, CRY1* and related clock components (Bhatwadekar et al., 2015; Na et al., 2009; Z. Wang et al., 2025; Zhou et al., 2021). In particular, miR-27b-3p oscillates in metabolic tissues, notably mouse liver, and directly represses BMAL1 by binding its 31UTR, with functional consequences for BMAL1 protein rhythms and hepatic metabolism (Ma, Yin, Zhong, Liang, & Guo, 2019; W. Zhang et al., 2016). Human simulated night-shift studies further demonstrate that circadian misalignment alters circulating exosomal miRNA profiles, and that these exosomes are sufficient to reduce BMAL1 function and impair insulin sensitivity in naïve adipocytes, establishing a causal link between exosomal miRNAs, clock disruption, and metabolic dysfunction (Khalyfa et al., 2020; Khalyfa et al., 2017).

A further consideration, largely absent from the cachexia literature, is that the cargo loaded into sEVs is itself temporally regulated. Emerging evidence indicates that sEV release and miRNA composition vary with time of day, raising the possibility that tumor sEVs transmit not only molecular but also temporal information to recipient tissues (Church, Kadukhina, Aiello, & Finkielstein, 2025). Whether this holds for cancer-bearing organisms remains to be tested.

The present work therefore tests the hypothesis that tumor-derived sEVs, by delivering rhythmically packaged miRNA cargo, contribute to the circadian disruption observed in host tissues of cancer-bearing organisms. This hypothesis is consistent with the causal link between EV miRNAs and clock-gene regulation established in non-cancer settings. Accordingly, we demonstrate that the PDAC secretome, and the sEVs within it, are sufficient to disrupt the circadian clock and induce myotube atrophy. We characterize the temporal dynamics of PANC-1 sEV release and of their miRNA cargo, and we resolve the cargo into two functionally distinct pools: a rhythmically packaged subset, whose secretion is phase-coordinated and which carries time-of-day information to the recipient tissue; and a constitutively secreted subset that is non-rhythmic at the source but is nonetheless capable of perturbing the recipient circadian clock once delivered. Individual miRNAs from both pools exert distinct and non-redundant effects on circadian period, myotube diameter, and mitochondrial bioenergetics, indicating that recipient-tissue clock disruption can arise from rhythmic as well as from tonic sEV miRNA input, reshaping muscle metabolism along three divergent trajectories; energetic, high-metabolic, and glycolytic. We next intersect the tumor sEV secretome with the serum sEV miRNA content of PDAC patients and with the pancreatic cancer miRNA-Seq landscape across 495 tumors. This analysis defines a stable, broadly tumor-abundant, patient-detectable miRNA signature that converges on circadian, proteostatic, and cachexia-relevant gene networks. Collectively, these findings position PDAC-derived sEV miRNAs as integrated, temporally organized mediators of the circadian–cachexia axis and as a previously uncharacterized class of systemic signals linking tumor biology and the clock to peripheral tissue dysfunction.

## Materials And Methods

### Cell Culture

The human pancreatic cancer cell line PANC-1, the murine fibroblast cell line NIH3T3, and the murine myoblast cell line C2C12 were purchased from American Type Culture Collection (ATCC, Manassas, VA). U2OS BMAL1:Luc cells were generously provided as a kind gift by Dr. John B. Hogenesch. PANC-1, NIH3T3, and C2C12 cells were maintained in DMEM supplemented with 10% fetal bovine serum and 0.5% Penicillin-Streptomycin. HPDE cells were maintained in Keratinocyte Serum-Free Media (KSFM, Invitrogen) supplemented with KSFM supplements, including epidermal growth factor (EGF) and bovine pituitary extract (BPE) (Invitrogen). All cell lines were maintained at 37 °C in a humidified atmosphere containing 5% CO_2_.

### Real Time Bioluminescence Recording

Real time bioluminescence recording was used to measure circadian periodicity in NIH3T3 *Bmal1*:Luc cells (Addgene, #68833). NIH3T3 *Bmal1*:Luc cells were seeded at 3×10^5^ in 35mm plates and allowed to adhere overnight (> 18 h). Following adherence, cells were incubated for 1 h in DMEM supplemented 200nM dexamethasone (MP Biomedicals) to synchronize the circadian clock. Cells were then washed with 1xPBS, and 2mL recording media was added. For conditioned media experiments, 2mL recording media (DMEM, Phenol Red Free, 2% FBS, 4mM L-Glutamine, 1mM Sodium Pyruvate, 10mM HEPES) for control, or 2mL conditioned media supplemented with the same components, was added immediately prior to recording. To test the effect of sEVs on circadian periodicity, cells were washed with 1xPBS and incubated with 2mL of recording media (DMEM Phenol Red Free, 2% FBS, 2mM L-Glutamine, 1mM Sodium Pyruvate, 10mM HEPES). sEV suspensions or PBS (control) were added to the respective plates immediately prior to recording. Real time bioluminescence recording was performed using either AB-2550 Kronos Dio (ATTO) or the LumiCycle 32-channel automated luminometer (Actimetrics). Bioluminescence was continuously recorded for at least 5 days. In all cases, raw data were collected at the end of dexamethasone synchronization (t=0) and throughout the duration of the experiment. For circadian period analysis, data starting at t=24 h post-synchronization were used. Period length was calculated using the BioDare2 software (biodare2.ed.ac.uk, University of Edinburgh) with either the fast Fourier transform-nonlinear least squares (FFT-NLSS) or MFourFit algorithm.

### Conditioned Media Generation and EV Isolation

PANC-1 cells were seeded at 5×10^5^ in a 100-mm tissue culture-treated plate and cultured until 60-80% confluency. PANC-1 cells were washed twice with serum-free DMEM before incubation with either serum-free DMEM or serum-free DMEM without phenol-red (for live bioluminescence imaging) for 72 h to generate conditioned media.

Small EVs (sEVs) were isolated using ExoQuick-TC exosome precipitation reagent (Systems Biosciences) according to manufacturer’s instructions. Briefly, conditioned media samples prepared as described above were centrifuged at 3000 xg for 15 min at 4° C to remove cells and cell debris. Supernatants were transferred to clean 15 mL tubes, ExoQuick-TC reagent was added at a 1:5 ratio (reagent:conditioned media), and tubes were rotated at 4° C to mix, then left upright overnight (> 18 h). After incubation, samples were centrifuged at 1500 xg for 30 min at 4° C, the supernatant was removed, and a second centrifugation at 1500 xg for 5 min at 4° C removed residual supernatant. sEVs were resuspended in 1x PBS or NP-40 lysis buffer (0.5% NP-40, 10mM Tris-HCl, 137mM NaCl, 1mM EDTA, 10% glycerol, with phosphatase and protease inhibitors) and stored at −80° C until use.

### Real Time Quantitative PCR (RT-qPCR)

Cells were harvested using TRIzol (ThermoFisher) and stored at −80° C until used. Total RNA was isolated using Direct-zol RNA Miniprep Kit (Zymo Research) with Zymo-Spin IICR columns, following manufacturer’s instructions. Total RNA concentration and quality were determined by spectrophotometry using a Nanodrop ND-2000 (Thermo Fisher Scientific). For cDNA synthesis, 1µg total RNA was used with the LunaScript RT SuperMix Kit (New England Biolabs), according to manufacturer guidelines. RT-qPCR was performed using the PowerUp SYBR Green Master Mix (Applied Biosystems) on a CFX Opus 384 Real-Time PCR System (Bio-Rad). Relative mRNAs expression was determined using the ΔΔCt method, with *Tbp* used as the reference gene for all analyses.

### Western Blotting

Western blotting was used for the determination of relative protein expression. Proteins were resolved by SDS-PAGE prior to semi-dry transfer to PVDF membranes. Membranes were blocked with 5% skim milk in TBST for 1 h prior to probing with primary antibodies diluted in TBST/skim milk, followed by HRP-conjugated secondary antibodies and detection with West Pico chemiluminescent solution (Thermo Fisher). Bio-Rad ImageLab software was used for image acquisition. For sEV characterization, PANC-1 cells and isolated sEV samples were lysed in ice-cold NP-40 lysis buffer; lysates were centrifuged at 14,000 ×g for 10 min at 4 °C, and supernatants were stored at –80 °C. Protein abundance was determined using the Pierce BCA protein assay kit (Thermo Scientific). Lysates (15 µg) were reduced in Laemmli sample buffer and separated on 12.5% SDS-PAGE. Proteins were transferred to PVDF membranes and blocked with 5% skim milk in TBST. Membranes were incubated overnight at 4 °C with the following primary antibodies: CD9 (Santa Cruz, sc-13118), TSG101 (Bethyl, A303-506), Alix (Abcam, ab117600), c-Myc (Abcam, ab32072), Calnexin (Cell Signaling, 2679), and ApoA2 (Invitrogen, PA5-114863). Membranes were washed with TBST and incubated for 1 h at room temperature with appropriate secondary antibodies.

### Nanoparticle Tracking Analysis

Nanoparticle tracking analysis (NTA) was utilized for nanoparticle quantification and size estimation. Samples were analyzed using a NanoSight NS300 running NTA v3.4 software (Malvern, UK), equipped with a 405 nm blue laser. Immediately prior to measurement, sEV suspensions were briefly vortexed at high speed and sonicated for 30 sec in a water bath to disperse aggregates. Nanoparticles were captured for 60 sec at 20° to 22°C, with viscosity set to that of water (0.915-0.918 cP), camera level 15, detection level set variably, and all other parameters at default. For scatter mode NTA and fluorescent NTA (fNTA) performed by AlphaNanoTech LLC, samples were analyzed on a Zetaview Quatt (Particle Metrix) according to AlphaNanoTech’s established protocols.

### Electron Microscopy

Negative stain electron microscopy was performed on isolated sEV samples using FCF200-CU formvar grids (Electron Microscopy Sciences). Grids were glow-discharged using a Pelco easiGlow system (TED PELLA), after which 10μL of sEVs resuspended in PBS or HEPES buffer was applied. Samples were dried prior to counter-staining with 10μL UranyLess EM stain (Electron Microscopy Sciences). EM was performed on an FEI Tecnai G2 Spirit Twin Transmission Electron Microscopy (ThermoFisher) equipped with Gatan Rio CCD camera.

### Calcein-AM Staining of EVs

Membrane integrity of isolated PANC-1 sEVs was assessed by Calcein acetoxymethyl-ester (Calcein-AM) staining. sEV suspensions in PBS were incubated with 10 µM Calcein-AM (Invitrogen, C3100MP) for 30 min at 37 °C, protected from light. Stained sEVs were imaged on a confocal fluorescence microscope, and intact vesicles were identified by the appearance of discrete fluorescent puncta corresponding to intracellular hydrolysis of the membrane-permeable substrate.

### Myoblast Differentiation

C2C12 myoblasts were seeded at 3×10^5^ per well in 6-well culture-treated plates and allowed to adhere overnight (>18 h). The culture medium was replaced with DMEM, 2% horse serum, 0.5% Penicillin-Streptomycin to induce differentiation into mature myotubes. Cells were cultured at 37 °C in a humidified atmosphere containing 5% CO_2_ with medium changed every 2 days over a minimum of 5 days.

### Transwell Co-Culture System

PANC-1 and NIH3T3 cells were seeded at 1×10^3^ per Transwell inserts (8-μm pore size; Corning) and cultured until reaching 60-80% confluency. The inserts were then transferred onto 6-well treated plates containing differentiated C2C12 myotubes and co-cultured for 48 h. As a positive control for atrophy induction, 100 μM dexamethasone was added. Images were captured every 12 h, with baseline measurements taken at 0 h.

### Imaging And Myotube Measurements

Myotube size was assessed by measuring myotube diameter. Digital images were acquired at 20x magnification on a Leica DMi1 inverted microscope equipped with a FLEXACAM C1 camera (Leica Microsystems). Average diameters were calculated from a minimum of n=10 randomly selected fields per well and n=3 replicate wells per time-point using ImageJ (version 2.16.0/1.54). Myotube diameter was defined as the average of three measurements taken at evenly spaced points along the length of each myotube.

### Time-Course Small EV Collection And Isolation

PANC-1 cells were seeded at 5×10^5^ in a 100 mm tissue culture-treated plates and cultured until 60-80% confluency. Cells were washed twice with serum-free DMEM and then incubated with serum-free DMEM for 24 h to limit serum-starvation artefacts. Cells were synchronized with 200nM dexamethasone for 1 h, the synchronization medium was replaced with fresh serum-free DMEM, and conditioned media were collected from the same 100 mm dish (N=3) every 4 h for the duration of the time-course, with plates washed once with serum-free DMEM between time-points to remove carry-over from the preceding 4 h interval. sEVs were isolated using ExoQuick-TC, as described above. Isolated sEV samples were stored at −80° C until shipment to third-party laboratory for small RNA-seq.

### EV Treatment In C2C12 Myotubes

sEVs from PANC-1 cells were generated as previously described. Differentiated C2C12 myotubes were treated with sEVs, PBS, or 100 μM dexamethasone and cultured for 72 h. Images were captured every 24 h, with baseline measurements taken at 0 h.

### Patients Serum Samples

Serum samples were provided by the Englander Institute for Precision Medicine at Weill Cornell Medicine. All procedures were conducted in accordance with institutional ethical guidelines and under Virginia Tech IRB protocol IRB-24-022. Six patients with pancreatic cancer (n=4 pancreatic adenocarcinoma, n=1 neuroendocrine pancreatic tumor, n=1 lymphohistiocytoid malignant mesothelioma of the pancreas). Clinical data including diagnosis, age at diagnosis, sex, prior treatment lines, and comorbidities were collected (Fig. 5A). Patients ranged in age from 46 to 77 years and had received between 1 and 7 prior lines of treatment. Small EVs were isolated using ExoQuick ULTRA EV Isolation Kit for Serum and Plasma (Systems Biosciences) according to manufacturer’s instructions.

### Small RNA Seq Analysis

Isolated sEVs from PANC-1 cells and serum patients were sent for small RNA-Seq analysis at Norgen Biotek (Ontario, CA). RNA isolation, quality control, small RNA-seq, adaptor trimming, filtering, and mapping were all performed by Norgen Biotek (workflow at https://norgenbiotek.com/services/small-rna-sequencing). RNA was isolated using Norgen’s Exosomal RNA Isolation Kit (Cat. 58000). Small RNA-seq was performed on an Illumina NextSeq 550 System with the NextSeq 500/550 High Output Kit v2 (51 Cycles using a 75-Cycle Kit). Library preparation used Norgen’s Small RNA Library Prep Kit for Illumina (Cat. 63600). Raw reads were mapped against miRBase v21 (miRNA), gtRNAdb (tRNA), RNAdb (piRNA), and Gencode v21/hg38 (genome).

### Mirna Sequencing And Quantification

For sEVs, raw sequencing reads were processed by Norgen Biotek. miRNA read counts and counts per million (CPM) were provided in the Norgen Service Report For patient serum samples, raw FASTQ files were uploaded to the Galaxy bioinformatics platform (https://usegalaxy.org/). Sequences were trimmed with the “Trim Galore!” tool to remove Illumina small RNA adapters, then collapsed with the miRDeep2 Quantifier using miRbase v21 precursor and mature sequences as reference. Normalized read counts (CPM) were obtained from miRDeep2 for each replicate of each patient sample (n=2 replicants per patient, n=6 patients), and replicates were averaged to yield a per-patient mean CPM.

### EV Expression Ranking And GO/KEGG Enrichment

miRNAs were ranked by mean log2 CPM expression across all nine EV time-points and the top 35 most expressed miRNAs were identified. Gene Ontology (GO) Biological Process enrichment was performed for validated targets of 11 out of these top 35 miRNAs. Experimentally validated miRNA-target interactions were retrieved from miRTarBase using the multiMiR R package. Target gene symbols were converted to Entrez IDs with bitr() from clusterProfiler. GO-BP enrichment used enrichGO() (OrgDb = org.Hs.eg.db, ontology = BP, BH correction, p<0.05, q<0.2). The top 35 terms by adjusted p-value were retained for visualization. Results were displayed as dot plots and per-miRNA contribution heatmaps.

### miRNA Transfection

Dharmacon miRIDIAN miRNA mimics and negative control mimics were purchased from Horizon Discovery (Cambridge, UK). Cell transfection used Lipofectamine RNAiMax Transfection Reagent (ThermoFisher) according to manufacturer’s guidelines. miRNA mimic stock solutions (20 μM) were diluted to the desired concentration in Opti-Mem reduced-serum medium (ThermoFisher) and mixed with RNAiMax at 0.75 μl/cm^2^ immediately prior to transfection. Cells were transfected with miRNA-RNAiMax complexes for 24 h.

Sequences of all miRNA mimics and miRNA mimic negative controls are: Negative control: UCACAACCUCCUAGAAAGAGUAGA (MIMAT0000039); hsa-miR-27b-3p: UUCACAGUGGCUAAGUUCUGC (MIMAT0000419); hsa-miR-99b-5p: CACCCGUAGAACCGACCUUGCG (MIMAT0000689); hsa-miR-127-3p: UCGGAUCCGUCUGAGCUUGGCU (MIMAT0000446); hsa-miR-191-5p: CAACGGAAUCCCAAAAGCAGCUG (MIMAT0000440); hsa-miR-615-3p: UCCGAGCCUGGGUCUCCCUCUU (MIMAT0003283); hsa-miR-10a-5p: UACCCUGUAGAUCCGAAUUUGUG (MIMAT0000253); hsa-miR-183-5p: UAUGGCACUGGUAGAAUUCACU (MIMAT0000261); hsa-miR-26a-5p: UUCAAGUAAUCCAGGAUAGGCU (MIMAT0000082); hsa-miR-30c-5p: UGUAAACAUCCUACACUCUCAGC (MIMAT0000244); hsa-miR-92a-3p: UAUUGCACUUGUCCCGGCCUGU (MIMAT0000092); hsa-let-7f-5p: UGAGGUAGUAGAUUGUAUAGUU (MIMAT0000067).

### Live Bioluminescence Recording In Transfected Cells

NIH3T3 *Bmal1*:Luc cells were seeded at 1.5×10^5^ per 35 mm plate and allowed to adhere overnight (>18 h). After adhesion, cells were transfected with 25 nM miRNA mimic or negative control for 24 h. Following transfection, the medium was replaced with 2 ml of DMEM supplemented with 10% FBS and 0.5% Penicillin-Streptomycin and synchronized with 200 nM dexamethasone for 1 h. Cells were then washed with PBS and 2 ml recording media (DMEM Phenol Red Free, 2% FBS, 4 mM L-Glutamine, 1 mM Sodium Pyruvate, 10 mM HEPES) was added immediately prior to recording. Live bioluminescence recording was performed as described above.

### Myotube Transfection and Measurement

C2C12 myoblasts were seeded at 3×10^5^ in 6-well plates and allowed to adhere overnight (> 18 h). The medium was replaced with differentiation medium and cells were differentiated for 5 days in DM. Mature myotubes were transfected with 25 nM miRNA mimic or negative transfection control for 24 h. Following transfection, the medium was replaced with differentiation media (DMEM supplemented with 2% horse serum, 0.5% Penicillin-Streptomycin). For positive control wells, 100 μM dexamethasone was added immediately prior to imaging (T=0) to induce myotube atrophy. Images were taken every 24 h for 48 h, with baseline measurements at 0 h.

### Seahorse XF Metabolic Phenotyping

C2C12 cells were transfected with individual miRNA, tested in two independent experiments and subjected to Seahorse XF analysis 48 h post-transfection. Oxygen consumption rate (OCR) and extracellular acidification rate (ECAR) were measured at baseline and following sequential injection of oligomycin (1 µM), FCCP (1 µM), and rotenone/antimycin A (0.5 µM each). Measurements were normalized to protein concentration for each well. Metabolic phenotypes were assigned based on resting and maximal OCR/ECAR profiles relative to negative-control transfected cells. Data are presented as mean ± SEM.

### Assessment of miRNA Expression and Abundance Across Pancreatic Cancer Samples

miRNA expression data for pancreatic cancer samples were obtained from the NIH-GDC data portal (https://portal.gdc.cancer.gov/), filtering by “primary site: pancreas; experimental strategy: miRNA-Seq; data type: miRNA Expression Quantification.” The resulting selection comprised 495 tumor samples quantified as reads per million (RPM), drawn from the TCGA-PAAD, CPTAC-3, and HCMI-CMDC. Because the dataset reports miRNA expression at the precursor (hairpin) level rather than strand-specific mature level, each mature miRNA was mapped to its corresponding precursor(s); for miRNAs with multiple genomic precursors, expression values were aggregated.

To assess relative expression magnitude, miRNA abundance was evaluated within each sample by ranking all detected miRNAs according to RPM. A miRNA of interest was considered highly expressed in a given sample when any of its corresponding precursor was among the top 50 most abundant miRNAs in that sample. The frequency with which each miRNA met this criterion across the cohort was calculated and expressed as both absolute counts and percentages of samples.

### Hierarchical Clustering Of Patient Mirna Profiles

Unsupervised hierarchical clustering of patient serum miRNA expression profiles was performed using Euclidean distance and Ward’s linkage method, implemented with hclust() in R. Three clusters were identified and manually assigned: Cluster 1 (P1, P3, P5), Cluster 2 (P4), and Cluster 3 (P2, P6) (Fig. 5A).

### Overlap Analysis: Patient Serum vs Tumor EV miRNAs

Each of the 428 patient miRNAs was assessed for detection in the EV dataset at ≥1 of 9 time-points. A total of 359 miRNAs were shared between patient serum and tumor EVs. The overlap was visualized as a Venn diagram and an UpSet plot stratified by patient (Fig. 5A, B). A stability filter requiring detection at all 9/9 EV time-points was applied to the EV dataset, yielding 494 constitutively secreted miRNAs. Intersection with the 428 patient miRNAs produced 42 stable candidate miRNAs present in both patient serum and stably secreted in tumor sEVs across all timepoints.

### Statistical Analysis and Visualization

All statistical analyses were performed in R (v4.5.2) and Graphpad Prism. Rhythmically expressed sEV miRNA were detected using the R package MetaCycle v1.2.0 with default settings (https://github.com/gangwug/MetaCycle). Differential expression used DESeq2 (v1.50.2) with Benjamini-Hochberg (BH) correction. GO enrichment analyses used clusterProfiler (v4.12) with BH correction. For MetaCycle, we used the meta2d function with a minimum period of 18 h and maximum period of 28 h (default period = 24 h). meta2d combines Lomb-Scargle, Jonckheere-Terpstra-Kendall (JTK), and ARSER (ARS) algorithms; p values from each were combined by Fisher’s method, with p<0.05 considered rhythmic. Visualizations were generated with ggplot2 (v4.0.2) and ggrepel (v0.9.8). Rayleigh R statistics were computed in R using the circular package against a null hypothesis of uniform phase distribution. All analyses used human miRNA annotations from miRBase v22.

## Results

### PDAC Secretome Drives Conserved, Multi-Level Circadian Disruption Across Cell Types

Consistent with the cell-intrinsic clock attenuation reported across pancreatic ductal adenocarcinoma (Jiang et al., 2016; Schwartz et al., 2023), MetaCycle analysis of dexamethasone-synchronized PANC-1 cells failed to detect statistically significant rhythmicity for any of the five core clock transcripts examined—BMAL1, PER2, CRY2, NR1D1, and DBP—over a 36 h time-course (Suppl. Fig. 1). The absence of a robust autonomous oscillator in the tumor cell itself motivates a reciprocal experimental design, in which the cell-extrinsic action of the PDAC secretome is interrogated in recipient reporter cells whose clocks are intact. To determine whether the human pancreatic ductal adenocarcinoma (PDAC) secretome is capable of altering the circadian clock, we employed a real-time bioluminescence reporter system in two independent cell lines. U2OS *BMAL1*:Luc cells were treated with PANC-1 conditioned media (CM) at increasing concentrations (12.5%, 25%, 50%, and 100% of the recording media) and circadian periodicity was monitored continuously for at least five days (Fig. 1A, B). PANC-1 CM produced a dose-dependent shortening of the circadian period (one-way ANOVA, p=0.0009; Fig. 1B). Rayleigh-plot analysis further revealed that, in U2OS cells, CM at 12.5% and 25% concentrations significantly altered the phase of *BMAL1*:Luc oscillations (p=0.0050 and p=0.0007, respectively; Fig. 1C), whereas 25% and 50% CM additionally increased oscillation amplitude (p=0.0023 and p=0.0015; Fig. 1C). To assess whether this effect extends to non-transformed murine fibroblasts, the same experiment was performed in NIH3T3 *Bmal1*:Luc cells. PANC-1 CM similarly induced a dose-dependent shortening of the *Bmal1*:Luc period in NIH3T3 cells (one-way ANOVA, p=0.0087; Fig. 1D, E), demonstrating that the effect of PDAC-secreted factors on the circadian clock is conserved across species and cell types. In NIH3T3 cells, phase disruption was most pronounced at the highest CM concentration (100%, p=0.0032; Fig. 1F), while amplitude was significantly elevated across a broader concentration range (25–100%; Fig. 1F), indicating that the two cell reporter systems display distinct sensitivity thresholds for phase-versus amplitude-level circadian perturbation by the PDAC secretome.

**Figure 1.**
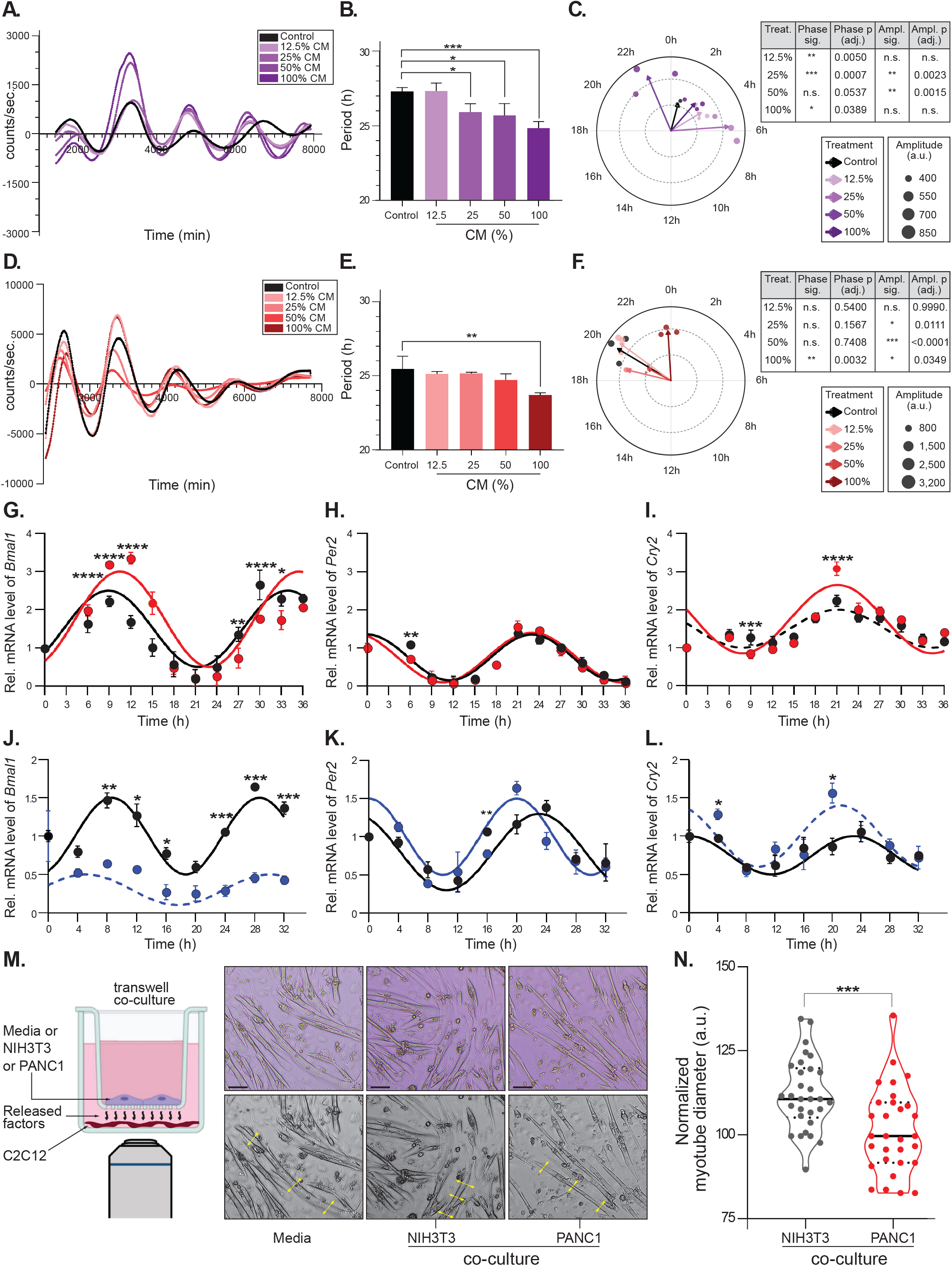
PDAC CM Promote Circadian Dysregulation. **(A)** Bioluminescence recording, (**B**) period analysis, and (**C**) phase and amplitude analysis of U2OS *BMAL1*:Luc reporter cells treated with PANC-1 CM at 12.5%, 25%, 50%, and 100% of the recording media. (**D**) Bioluminescence recording, (**E**) period analysis, and (**F**) phase and amplitude analysis of NIH3T3 *Bmal1*:Luc reporter cells treated with PANC-1 CM at the same concentrations. For all bioluminescence experiments, at least three complete oscillations were included in the period estimation, excluding the first 24 h. Mean ± SD of relative mRNA expression of NIH3T3 core clock genes *Bmal1* (**G**), *Per2* (**H**), and *Cry2* (**I**) measured over 36 h in response to PANC-1 CM. Relative mRNA levels of core clock genes in synchronized C2C12 myotubes over 32 h following treatment with PANC-1 CM: (**J**) *Bmal1*, (**K**) *Per2*, and (**L**) *Cry2*. Cosine curves were fit for visualization purposes only; solid lines represent rhythmic oscillations (p<0.05) detected by MetaCycle, while dashed lines indicate loss of statistical rhythmicity (Suppl. Table 1). (**G–L**) Black: Control; (**G–I**) Red: PANC-1 CM; (**J–L**) Blue: PANC-1 CM. (**M**) Schematic representation and representative images of mature C2C12 myotube atrophy in response to NIH3T3 or PANC-1 released factors using a Transwell co-culture system; three measurements per myotube (yellow arrows) were used to quantify shortening. (**N**) Quantification of normalized myotube diameter under NIH3T3 vs PANC-1 co-culture, normalized to NIH3T3 co-culture control. One-way ANOVA: (**B**) p=0.0009, (**E**) p=0.0087. (**B, E**) Dunnett’s post-hoc test: *p<0.05; **p<0.01; ***p<0.001. (N) Student’s t-test: ***p<0.001.

To further dissect the molecular basis of CM-mediated circadian dysregulation, RT-qPCR was used to measure core clock transcripts in NIH3T3 fibroblasts treated with PANC-1 CM, with total RNA collected every 3 h over a 36-h period following circadian synchronization. PANC-1 CM (100%) lengthened the periodicity of *Bmal1* and *Per2* transcripts and produced a moderate phase delay (~2 h, p<0.001) in *Bmal1* (Fig. 1G–H). MetaCycle analysis confirmed that CM treatment altered the period, phase, and amplitude of core clock transcripts, with rhythmic oscillations represented as solid fitted cosine curves and loss of detectable rhythmicity as dashed lines (Fig. 1G–I, Suppl. Table 1). Of note, the bioluminescence reporter and RT-qPCR readouts captured the perturbation at distinct regulatory layers. The *Bmal1* promoter– driven luciferase reports the rate of transcriptional initiation, whereas RT-qPCR integrates transcription and mRNA turnover. The opposite directionality of the period change at these two layers therefore points to a post-transcriptional contribution to clock remodeling, a mechanism for which miRNAs constitute a parsimonious candidate, and one that we pursue in Figs. 3 and 4. Of additional interest, *Cry2* was non-rhythmic in control NIH3T3 cells but acquired statistically detectable rhythmicity upon CM treatment (Fig. 1I and Suppl. Table 1), underscoring the bidirectional nature of clock remodeling by tumor-secreted factors.

We next investigated whether PANC-1-derived secreted factors disrupt the circadian clock in skeletal muscle, a primary target of PDAC-associated cachexia. Mature C2C12 myotubes were treated with PANC-1 CM (100%) and total RNA was collected every 4 h over a 32-h period (Fig. 1J–L). CM treatment produced a complete ablation of detectable *Bmal1* rhythmicity in mature myotubes (Fig. 1J, Suppl. Table 1); *Per2* mRNA expression exhibited a shortened period (Fig. 1K, Suppl. Table 1), and *Cry2* similarly lost statistical rhythmicity (Fig. 1L, Suppl. Table 1). The effect of PANC-1 CM on the muscle clock was substantially more pronounced than in NIH3T3 fibroblasts, indicating cell-type-dependent sensitivity to PDAC-secreted factors and identifying skeletal muscle as a particularly vulnerable peripheral oscillator.

To determine whether PDAC-secreted factors also drive myotube atrophy, we employed a Transwell co-culture system that supports continuous paracrine communication without direct cell–cell contact. Mature C2C12 myotubes were co-cultured with either PANC-1 or NIH3T3 cells for 48 h (Fig. 1M). Co-culture with PANC-1 cells produced a significant reduction in myotube diameter relative to NIH3T3 co-culture (p<0.001; Fig. 1M, N), establishing that the PDAC paracrine secretome is, on its own, sufficient to induce myotube atrophy *in vitro*.

### PDAC-Derived sEVs are Sufficient To Promote Circadian Dysregulation and Myotube Atrophy

Having established that the PDAC secretome dysregulates the circadian clock and induces myotube atrophy, we asked whether small extracellular vesicles (sEVs) within that secretome are sufficient to mediate these effects. sEVs were isolated from PANC-1 conditioned media and characterized in accordance with established guidelines (Welsh et al., 2024) (Fig. 2A–D). Western blotting confirmed enrichment of canonical EV markers, including CD9 (a tetraspanin membrane protein) and TSG101 (a component of the ESCRT-I complex), with exclusion of the endoplasmic-reticulum marker Calnexin and the lipoprotein marker Apolipoprotein A2 (ApoA2) (Fig. 2A). Negative-stain electron microscopy revealed the cup-shaped morphology characteristic of sEV populations (Fig. 2B), and Calcein-AM staining confirmed intact vesicular membranes through the appearance of discrete fluorescent puncta (Fig. 2C). Fluorescent nanoparticle tracking analysis (fNTA) demonstrated a size distribution consistent with sEVs, with a mean diameter <200 nm (Fig. 2D).

**Figure 2.**
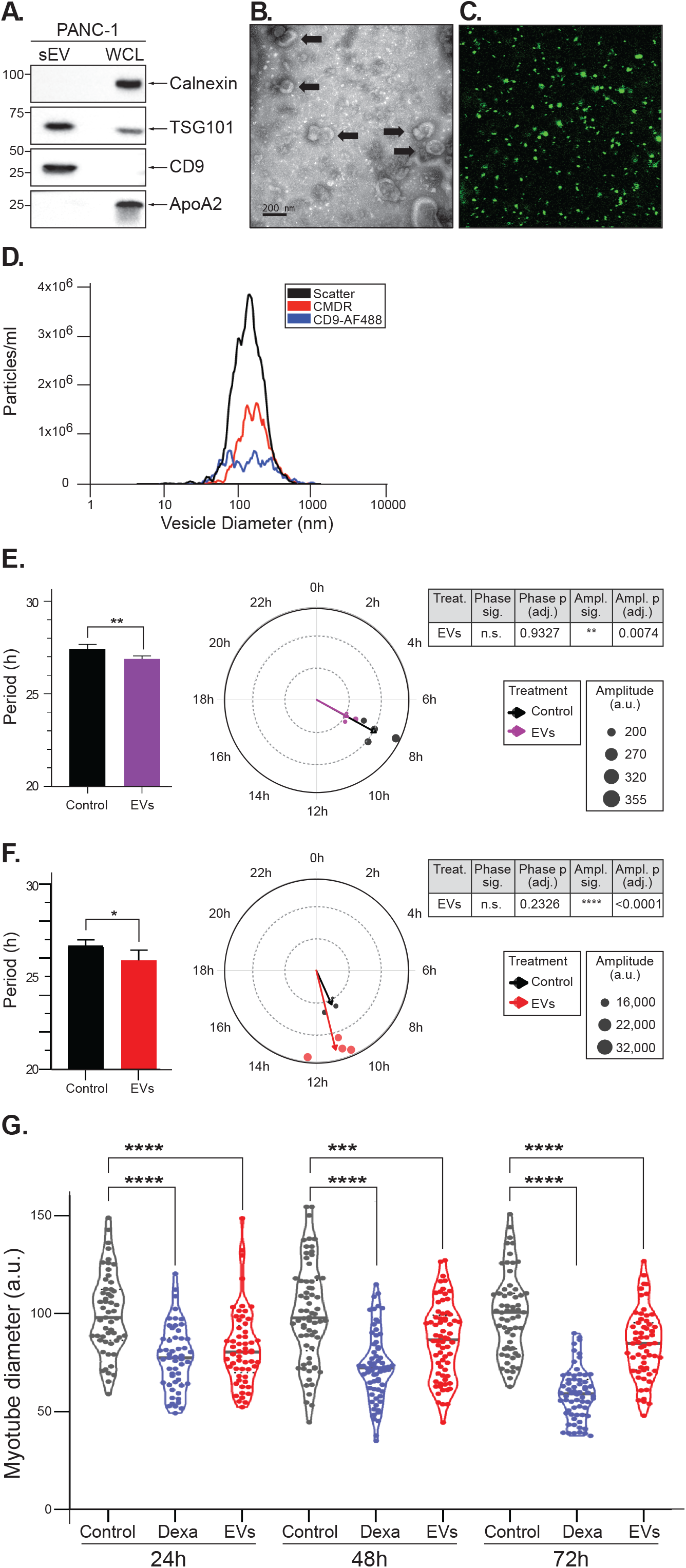
PDAC sEVs Promote Circadian Dysregulation and Myotube Atrophy. (**A**) Representative western blot for Calnexin, TSG101, CD9, and ApoA2 in PANC-1 whole-cell lysates (WCL) and isolated sEVs. (**B**) Negative-stain electron microscopy of PANC-1 sEVs resuspended in PBS, showing the characteristic cup-shaped morphology typical of EVs. Black arrows: representative sEVs displaying cup-shaped morphology. (**C**) Calcein acetoxymethyl-ester (Calcein-AM) staining of PANC-1 sEVs showing distinct fluorescent puncta indicative of intact vesicular membranes. (D) Representative fluorescent NTA (fNTA) of PANC-1 sEVs incubated with a fluorescently conjugated antibody against CD9. CMDR: CellMask Deep Red; Scatter: total particles; CD9-AF488: CD9-conjugated fluorescent-antibody-labeled particles. (**E**) Period, phase, and amplitude analyses of BMAL1:Luc oscillations in U2OS reporter cells treated with sEVs isolated from PANC-1 CM. (**F**) Period, phase, and amplitude analyses of Bmal1:Luc oscillations in NIH3T3 reporter cells treated with sEVs isolated from PANC-1 CM. For all bioluminescence experiments, at least three complete oscillations were included in the period estimation, excluding the first 24 h. Student’s t-test: (**E**) period p<0.01, amplitude p=0.0074; (F) period p<0.05, amplitude p<0.0001. (**G**) Myotube atrophy in response to PANC-1 sEVs over 72 h. Measurements were taken from at least 10 random fields in n=3 wells and normalized to control (without sEVs) for each time-point. Dexamethasone (Dexa) is included as a positive control for atrophy. ****p<0.0001; ***p<0.001.

**Figure 3.**
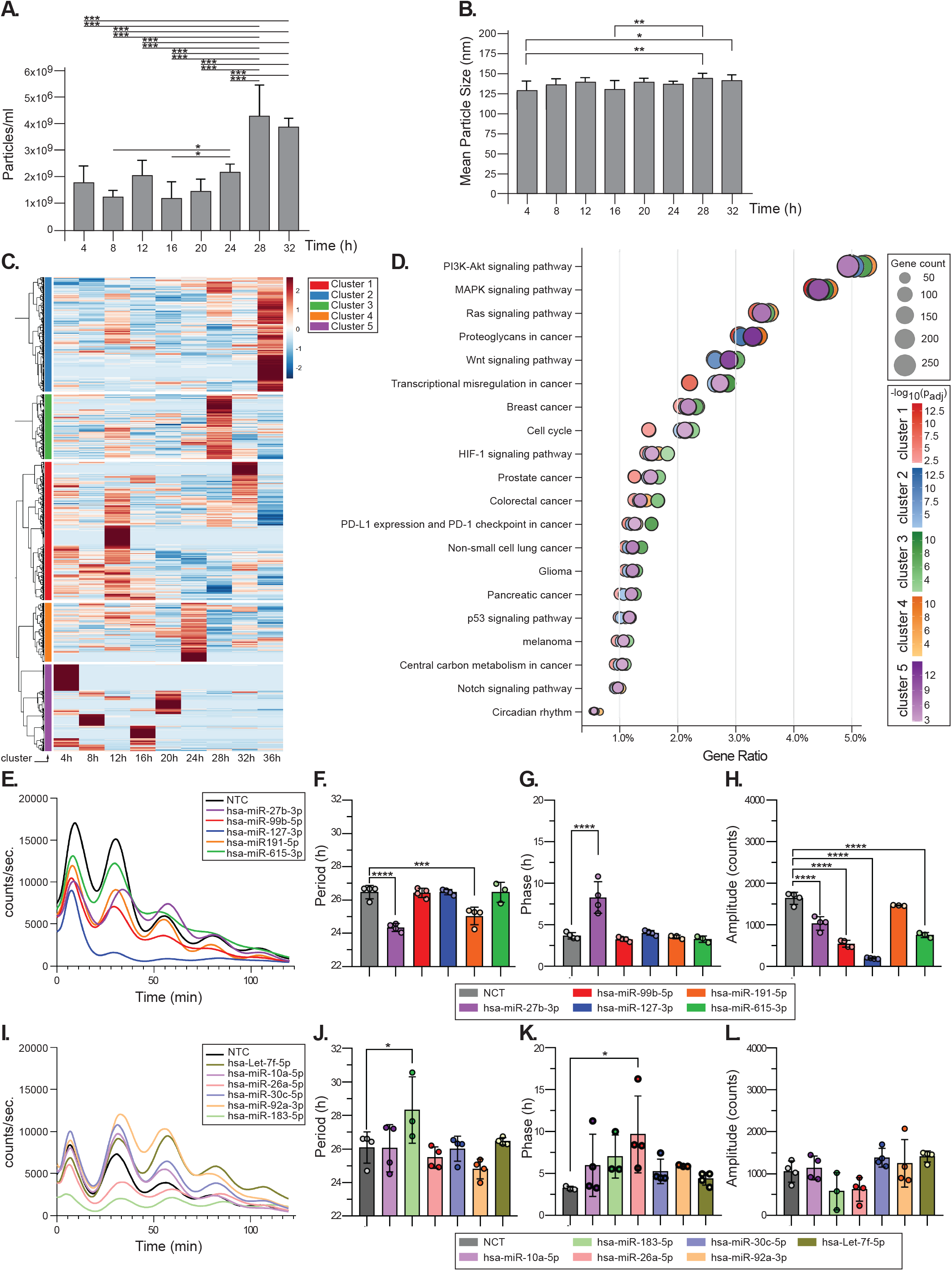
Time-Of-Day-Dependent Release and miRNA Cargo of PANC-1 sEVs. **(A)** Concentration of sEVs and (**B**) mean particle size of sEVs isolated from CM of circadian-synchronized PANC-1 cells collected every 4 h for 32 h (n=3 biological replicates per time-point). (**C**) Heatmap displaying z-score-normalized log2(CPM+1) expression of 1,426 miRNAs detected in PANC-1 sEVs collected at 9 consecutive time-points. Hierarchical clustering used Ward.D2 linkage with Euclidean distance. Rows represent individual miRNAs; columns represent time-points. Color scale: red = above-mean expression; blue = below-mean expression. The dendrogram on the left shows hierarchical relationships among miRNAs. Five clusters were identified with distinct temporal expression patterns: C1 (Red, n=352), C2 (Blue, n=200), C3 (Green, n=426), C4 (Orange, n=181), and C5 (Purple, n=267). The optimal number of clusters (k=5) was determined by combining the elbow method and silhouette analysis (mean silhouette score = 0.187). (**D**) Dot plots showing KEGG enrichment for miRNA clusters C1–C5. Target genes were predicted using miRDB (score ≥80) for each cluster’s miRNA set. The x-axis represents Gene Ratio; dot size represents the number of genes enriched in each pathway (Gene count). Dot color (C1: Red, C2: Blue, C3: Green, C4: Orange, C5: Purple) indicates statistical significance as −log10(padj); only the top 20 significant terms are shown. (**E**) Representative bioluminescence curves of synchronized NIH3T3 *Bmal1*:Luc cells transfected with 25 nM of miRNA mimics: hsa-miR-27b-3p, hsa-miR-99b-5p, hsa-miR-127-3p, hsa-miR-191-5p, hsa-miR-615-3p, or negative-transfection control (cel-miR-67). (**F**) Period, (**G**) phase, and (**H**) amplitude analyses of NIH3T3 *Bmal1*:Luc cells transfected with the miRNAs in panel E. (**I**) Representative bioluminescence curves of synchronized NIH3T3 *Bmal1*:Luc cells transfected with 25 nM of: hsa-let-7f-5p, hsa-miR-10a-5p, hsa-miR-26a-5p, hsa-miR-30c-5p, hsa-miR-92a-3p, hsa-miR-183-5p, or negative-transfection control (cel-miR-67). (**J**) Period, (**K**) phase, and (**L**) amplitude analyses of NIH3T3 *Bmal1*:Luc cells transfected with the miRNAs in panel **I.** All data are presented as mean ± SD. Circadian period, phase, and amplitude were calculated using the FFT-NLLS algorithm in BioDare2 (https://biodare2.ed.ac.uk/). Statistical analyses of period, phase, and amplitude were performed by one-way ANOVA with Dunnett’s post-hoc test in GraphPad Prism. *p<0.05; **p<0.01; ***p<0.001; ****p<0.0001; n.s., not significant.

**Figure 4.**
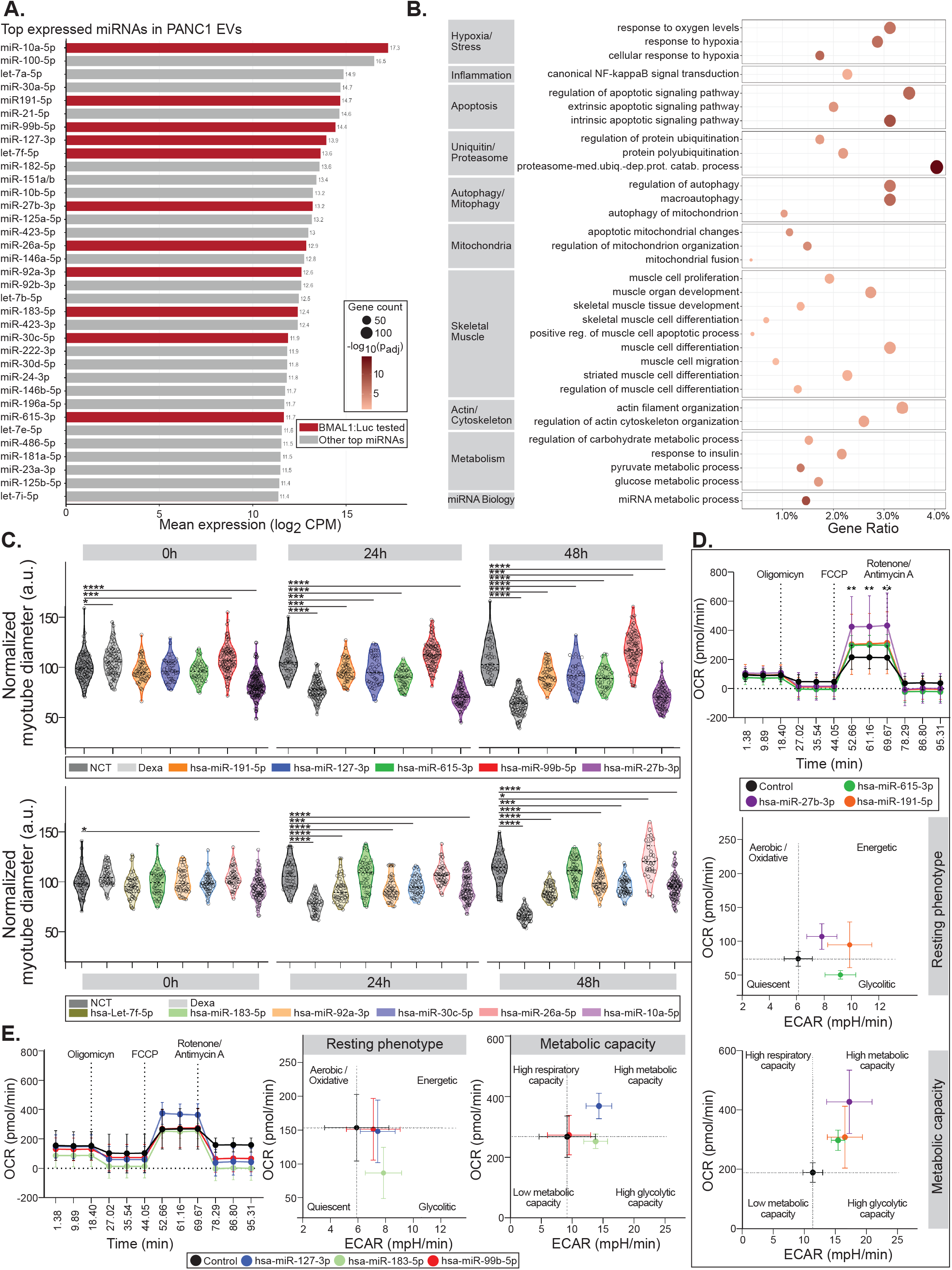
PDAC-Secreted miRNAs Induce C2C12 Myotube Atrophy and Remodel Bioenergetics. **(A)** Top 35 miRNAs by mean expression in PANC-1-derived sEVs. Bar plot of mean log2 CPM across 9 time-points (4–36 h). Red bars: miRNAs selected from the top 35 to be tested in the *BMAL1*:Luc reporter and atrophy assays; grey bars: remaining top-35 miRNAs. miRNAs are ranked in descending order of EV expression. (**B**) GO Biological Process enrichment of the experimentally validated targets (miRTarBase) of the 11 selected miRNAs. Terms are grouped into functional categories. Dot size represents the number of validated target genes associated with each term; dot color indicates Gene Ratio (proportion of input genes annotated to the term), from light pink (low) to dark red (high). Analysis performed with clusterProfiler. (**C**, top panel) Normalized C2C12 myotube diameter at 0, 24, and 48 h post-transfection with miR-27b-3p, miR-615-3p, miR-191-5p, miR-127-3p, miR-99b-5p, or negative-transfection control (NTC); dexamethasone (Dexa) included as positive control. (**C**, lower panel) Normalized C2C12 myotube diameter at the same time-points after transfection with hsa-let-7f-5p, miR-183-5p, miR-92a-3p, miR-30c-5p, miR-26a-5p, miR-10a-5p, NTC, or Dexa. (**D**) Oxygen consumption rate (OCR; top), resting-phenotype plot of basal OCR vs ECAR (middle), and metabolic-capacity plot of maximal OCR vs ECAR following FCCP (lower) for mature C2C12 myotubes 48 h post-transfection with miR-27b-3p, miR-615-3p, miR-191-5p, or NTC (Control). (**E**) Same panels for myotubes transfected with miR-127-3p, miR-99b-5p, miR-183-5p, or NTC. Sequential injections of oligomycin, FCCP, and rotenone/antimycin A were used to dissect mitochondrial respiration. Data are presented as mean ± SEM. (**C**) Measurements were taken from at least 5 random fields per well in N=3 wells; statistical analysis used 2-way ANOVA with Dunnett’s post-hoc correction: *p<0.05; **p<0.01; ***p<0.001; ****p<0.0001.

**Figure 5.**
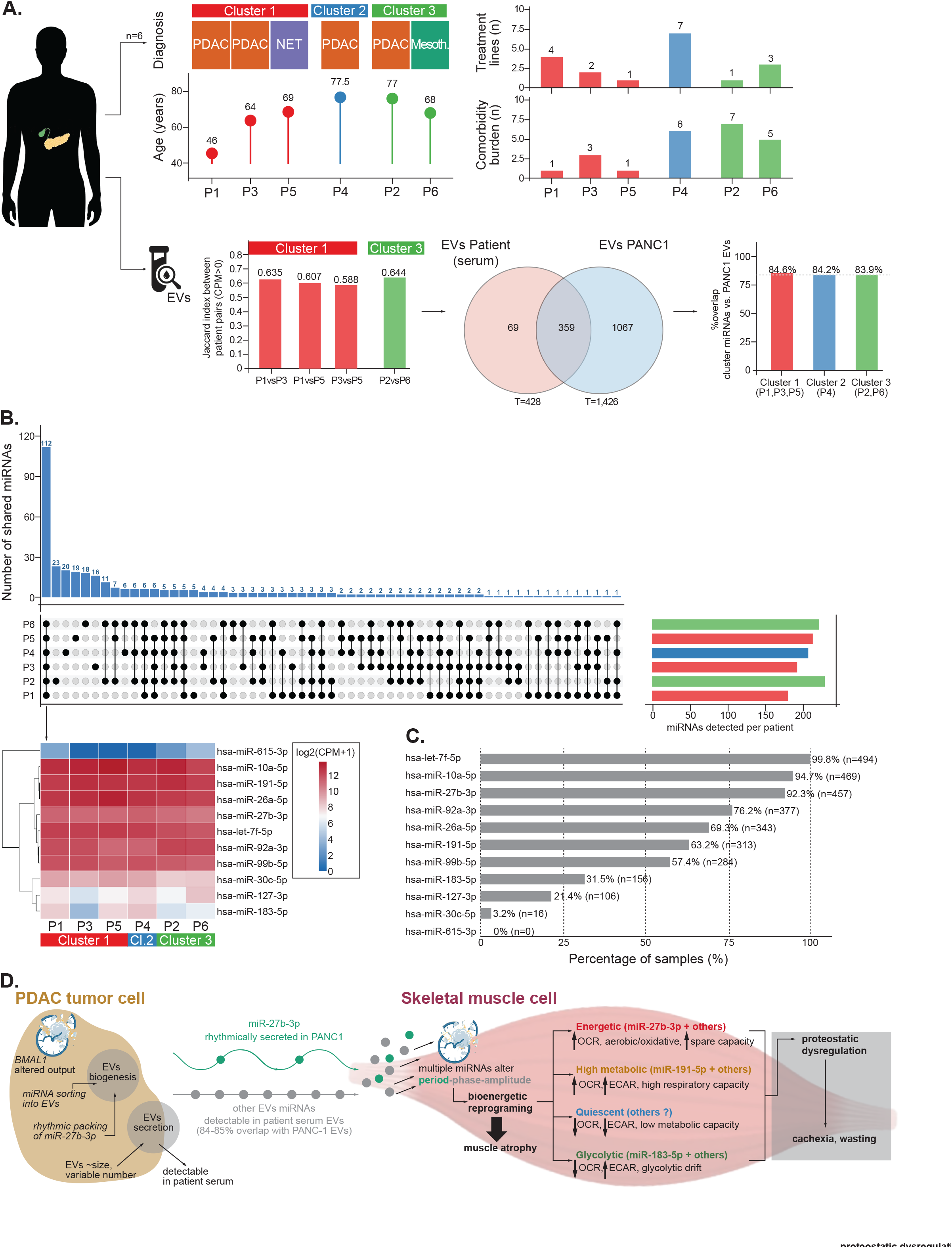
Patient-Circulating miRNAs Overlap with the Stable Secretome of Pancreatic Tumor-Derived Extracellular Vesicles. **(A)** Clinical and molecular characterization of six pancreatic cancer patients (n=6) stratified by unsupervised hierarchical clustering of serum sEV miRNA expression profiles. Three clusters were resolved: Cluster 1 (P1, P3, P5; red), Cluster 2 (P4; blue), and Cluster 3 (P2, P6; green). Diagnoses are indicated above each patient identifier: PDAC, pancreatic ductal adenocarcinoma; NET, neuroendocrine tumor; Mesoth., malignant mesothelioma of the pancreas. Clinical parameters shown include age at diagnosis (dot plot, years), number of prior treatment lines (bar chart, top right), and comorbidity burden (bar chart, bottom right). The Jaccard similarity index between within-cluster patient pairs (CPM>0) is plotted for Clusters 1 and 3, demonstrating high intra-cluster miRNA-profile similarity (range 0.588–0.644). A Venn diagram illustrates the overlap between miRNAs detected in patient serum sEVs (T=428) and in PANC-1 sEVs across 9 time-points (T=1,426); 359 miRNAs are shared between the two datasets. Bar charts indicate the percentage of cluster-specific patient miRNAs detectable in the PANC-1 sEV secretome: Cluster 1, 84.6%; Cluster 2, 84.2%; Cluster 3, 83.9%. (**B**) UpSet plot (top) displaying the combinatorial overlap of miRNAs detected in each patient’s serum sEVs. Vertical bars indicate the number of miRNAs shared by each intersecting patient combination defined in the connected dot matrix below; horizontal bars on the right show the total number of miRNAs detected per patient. Below the UpSet plot, a heatmap shows expression levels (log2(CPM+1)) of the 11 candidate miRNAs functionally validated in this study across all six patients. Rows are ordered by hierarchical clustering of miRNA expression profiles; columns are arranged by patient cluster assignment (Cluster 1, red; Cluster 2, blue; Cluster 3, green). Color scale ranges from blue (low expression) to dark red (high expression). (**C**) Frequency of the 11 candidate miRNAs among the top 50 most highly expressed miRNAs in pancreatic cancer tumor specimens. For each of 495 tumor samples from the GDC data portal (TCGA-PAAD, CPTAC-3, and HCMI-CMDC), miRNAs were ranked by RPM, and a given miRNA was classified as highly expressed if any of its corresponding precursors ranked within the top 50. Bars show the percentage of samples meeting this criterion; absolute sample counts are given in parentheses. (**D**) Proposed mechanistic model. Within the PDAC tumor cell, a dysregulated clock (BMAL1↓, altered rhythmic output) drives miRNA sorting into EVs during biogenesis; miR-27b-3p is uniquely (among those tested) and rhythmically packaged, while other sEV miRNAs are loaded constitutively. Secreted sEVs exhibit variable particle number with stable size, and these miRNAs are detectable in patient serum (84– 85% overlap with PANC-1 sEV miRNAs). In the recipient skeletal muscle cell, the full sEV miRNA cargo collectively disrupts the circadian clock, altering period, phase, and amplitude. In parallel, individual miRNAs reprogram bioenergetics along four non-redundant trajectories: Energetic (miR-27b-3p + others; ↑OCR, aerobic/oxidative, ↑spare respiratory capacity); High metabolic (miR-191-5p + others; ↑OCR, ↑ECAR, high respiratory capacity); Quiescent (others; ↓OCR, ↓ECAR, low metabolic capacity); and Glycolytic (miR-183-5p + others; ↓OCR, ↑ECAR, glycolytic drift). The integration of circadian disruption, bioenergetic reprogramming, and proteostatic dysregulation collectively drives muscle-cell atrophy *in vitro*; whether and how these molecularly distinct insults produce a systemic cachexia phenotype *in vivo* remains to be determined (gray box).

To assess the effect of isolated sEVs on the circadian clock, U2OS *BMAL1*:Luc cells were treated with PANC-1-derived sEVs. sEV treatment shortened the circadian period (p<0.01; Fig. 2E) and produced a marked amplitude reduction (p=0.0074; Fig. 2E), without a significant phase shift. To extend the observation to a second reporter cell line, NIH3T3 *Bmal1*:Luc cells were treated identically; treatment shortened the *Bmal1*:Luc period (p<0.05; Fig. 2F) and altered oscillation amplitude (p<0.0001; Fig. 2F), again without a significant phase shift. The divergence in amplitude effect magnitude between U2OS and NIH3T3 cells is consistent with the cell-type-dependent sensitivity to PDAC-secreted factors observed at the CM level (Fig. 1C, F) and is compatible with a contribution of post-transcriptional mechanisms downstream of clock-gene transcription to sEV-mediated circadian remodeling.

We next asked whether isolated PANC-1 sEVs are sufficient to induce myotube atrophy. Differentiated C2C12 myotubes were treated with PANC-1 sEVs or vehicle control, and myotube diameter was measured every 24 h over a 72 h period (Fig. 2G). PANC-1 sEVs produced a significant reduction in myotube diameter at every time-point assessed, with dexamethasone serving as a positive atrophy control (Menconi, Gonnella, Petkova, Lecker, & Hasselgren, 2008) (Fig. 2G). Together, these results establish that PDAC-derived sEVs are sufficient to recapitulate both arms of the secretome phenotype: circadian dysregulation in clock-reporter cells and atrophy in mature myotubes.

### Time-Of-Day-Dependent Variation in PANC-1 sEVs Release, miRNA Cargo, And Functional Effects on the Circadian Clock

Whether the time of day shapes sEV release and composition is of immediate significance for identifying sEV subpopulations that may drive distinct aspects of cancer progression and for the design of EV-based biomarker studies. To characterize the temporal dynamics of sEV release, sEVs were collected from circadian-synchronized PANC-1 cells every 4 h over a 32 h window and quantified by nanoparticle tracking analysis (Fig. 3A, B). sEV concentration varied across time-points, with later collection windows (24, 28, and 32 h post-synchronization) yielding significantly higher sEV concentrations than earlier ones (Fig. 3A). Mean vesicle size was largely invariant across the time-course, with only modest but statistically significant differences between specific time-points [4 h vs. 28 h (p<0.01) and 32 h (p<0.05); 16 h vs. 28 h (p<0.05); Fig. 3B]. MetaCycle analysis did not detect statistically significant rhythmic oscillation of either sEV concentration or mean vesicle size, indicating that the observed variation in sEV release does not conform to a canonical circadian pattern under these experimental conditions; rather, it reflects time-dependent kinetics of vesicle accumulation.

Given the established role of miRNAs as stable, EV-enriched regulators of post-transcriptional gene expression, with the capacity to coordinately modulate complex gene networks including the circadian machinery, we focused on the miRNA cargo of PANC-1 sEVs and its temporal variation. Small RNA sequencing was performed on sEVs collected every 4 h for 36 h (n=3 biological replicates per time-point), yielding 1,426 detected miRNAs across all time-points. Hierarchical clustering of z-score-normalized expression revealed five distinct temporal clusters (k=5; Fig. 3C). KEGG pathway enrichment of the predicted and validated targets of the cluster-constituent miRNAs (Fig. 3D) showed that all five clusters were significantly enriched for major oncogenic signaling pathways, including PI3K–Akt, MAPK, and Ras, as well as Wnt, HIF-1, and p53. Cancer-type-specific enrichment, including pancreatic cancer, was apparent predominantly in clusters 3 and 5. Of particular significance, the KEGG circadian rhythm pathway was uniquely and significantly enriched in all clusters (Fig. 3D). The convergence of an unbiased target-level enrichment analysis on the same biological process functionally probed by our bioluminescence reporter (Fig. 1, Fig. 2E–F) establishes the circadian clock as a cell-extrinsic target of the PDAC sEV miRNA payload, rather than an artefact of any single experimental readout. Cell cycle, PD-L1/PD-1 checkpoint, and central carbon metabolism pathways were additionally enriched across multiple clusters, linking the temporally organized sEV miRNA program to proliferative, immunomodulatory, and bioenergetic networks of direct relevance to PDAC biology. MetaCycle analysis identified 67 miRNAs that were rhythmically expressed in PANC-1 sEVs (p<0.05) and that were distributed across four phase-defined windows (Supp. Fig. 2A). The heatmap representation of these 67 rhythmic species (Supp. Fig. 2A) resolves the temporal architecture of the cargo at single-miRNA resolution. Rather than distributing uniformly across the day, rhythmic miRNAs segregate into discrete phase blocks, with the strongest acrophase enrichment observed in the early (0–8 h) and mid-day (8–16 h) windows, indicating that sEV packaging follows a phase-coordinated rather than a stochastic schedule. Of immediate relevance to the candidates pursued downstream, hsa-miR-27b-3p is one of the most highly significant rhythmic species in the dataset (p<0.01) and falls within the mid-day phase window, anchoring its identification in subsequent functional assays (Figs. 3–5) within an objectively defined rhythmic subprogram rather than within the bulk constitutive cargo. Rayleigh analysis of the acrophase distribution of these 67 rhythmic miRNAs revealed their temporal organization across the 24 h cycle (Supp. Fig. 2B). The mean Rayleigh vector (R=0.34) is non-trivial yet far from unity, formally establishing that rhythmic loading is phase-biased without collapsing onto a single time-point, a property that distinguishes a genuinely temporally structured cargo from one driven by a single dominant peak, and that is consistent with phase-staggered, multi-miRNA secretion from the tumor cell. Together, these findings demonstrate that the miRNA composition of PDAC-derived sEVs varies in a time-of-day-dependent manner, with a defined subset exhibiting statistically robust rhythmic expression.

To identify the most abundant miRNA species in PANC-1 sEVs, all detected miRNAs were ranked by mean log2 CPM expression across the nine time-points. The top 35 most expressed miRNAs, irrespective of cluster assignment, are presented in Fig. 4A. From these, 11 miRNAs were selected to represent the various expression clusters, hsa-miR-10a-5p, hsa-let-7f-5p, hsa-miR-27b-3p, hsa-miR-615-3p, hsa-miR-191-5p, hsa-miR-26a-5p, hsa-miR-92a-3p, hsa-miR-99b-5p, hsa-miR-127-3p, hsa-miR-30c-5p, and hsa-miR-183-5p, based on their high and constitutive abundance in the sEV secretome and the enrichment of their validated targets in circadian, proteostatic, and muscle-relevant pathways (Fig. 3D and Fig. 4B); their selection therefore integrates expression prominence with predicted functional relevance to the clock–cachexia axis under investigation.

To assess whether sEV-derived miRNAs are themselves capable of modulating the circadian clock, NIH3T3 *Bmal1*:Luc cells were transfected with 25 nM of each candidate miRNA mimic and real-time bioluminescence recording was performed (Fig. 3E–L). Among the candidates, hsa-miR-27b-3p produced a significant shortening of the *Bmal1*:Luc period (~2.18 h, p<0.0001; Fig. 3F) and a pronounced phase shift (p<0.0001; Fig. 3G). hsa-miR-191-5p similarly shortened the period (~1.18 h, p<0.001; Fig. 3F) but, unlike miR-27b-3p, did not alter phase or amplitude (Fig. 3G, H). In the second group of miRNAs tested, hsa-miR-183-5p was the only candidate to significantly lengthen the *Bmal1*:Luc period (p<0.05; Fig. 3J), accompanied by a significant phase alteration (p<0.05; Fig. 3K). These results demonstrate that individual sEV-derived miRNAs exert distinct, and in some cases opposing, effects on circadian clock parameters, underscoring the multi-input nature of miRNA-mediated circadian regulation.

### sEV-Derived miRNAs Target Metabolic And Muscle Pathways, Induce Myotube Atrophy, And Reprogram Cellular Metabolism

Having identified miR-27b-3p, miR-191-5p, and miR-183-5p as sEV-derived miRNAs capable of modulating core clock oscillations (Fig. 3), we next asked whether the broader set of 11 candidate miRNAs collectively targets on gene networks of pathophysiological relevance to PDAC-associated muscle wasting. Gene Ontology (GO) enrichment analysis was performed using clusterProfiler and enrichGO on the experimentally validated miRNA targets retrieved from miRTarBase.

Enriched terms partitioned into functionally coherent categories spanning hypoxia and stress response, NF-κB signaling, apoptosis, ubiquitin–proteasome and autophagy/mitophagy pathways, mitochondrial organization, skeletal muscle biology, actin cytoskeleton, and carbohydrate metabolism (Fig. 4B). Of particular relevance to cachexia, targets were significantly enriched for terms governing skeletal muscle cell differentiation, muscle organ development, and muscle tissue development, as well as for regulators of mitochondrial organization, apoptotic mitochondrial changes, and mitophagy. Metabolic terms, including pyruvate metabolic process, glucose metabolic process, and response to insulin, were likewise overrepresented. Collectively, this target landscape nominates the candidate miRNAs as coordinated modulators of the proteostatic, mitochondrial, and metabolic machineries that are dismantled during cancer-induced muscle wasting.

To determine whether these miRNAs are individually sufficient to drive the atrophic phenotype, mature C2C12 myotubes were transfected with nanolipid-encapsulated mimics of each of the 11 candidates, and myotube diameter was quantified at 0, 24, and 48 h post-transfection, with dexamethasone (DEX) as a positive control (Fig. 4C). As shown in Fig. 4C (top panel), miR-191-5p, miR-127-3p, miR-615-3p, miR-99b-5p, and miR-27b-3p each elicited a significant reduction in myotube diameter relative to non-targeting control (NTC), with the effect emerging as early as 24 h and becoming most pronounced at 48 h, with miR-27b-3p producing atrophy of a magnitude comparable to DEX. In the lower panel of Fig. 4C, let-7f-5p, miR-92a-3p, miR-30c-5p, miR-26a-5p, and miR-10a-5p similarly reduced myotube diameter at 48 h. The near-universal atrophic response across chemically and seed-sequence-distinct miRNAs indicates that the PDAC sEV miRNA payload functions not as a single dominant effector but as a multi-miRNA program targeting the muscle proteome from multiple angles, an organization consistent with the GO enrichment signature in Fig. 4B.

We next tested whether the three miRNAs that exerted the strongest circadian effects, miR-27b-3p, miR-191-5p, and miR-183-5p (Fig. 3F, J), also reprogram bioenergetics, and whether they do so in a coherent manner. Mature C2C12 myotubes were transfected with the respective mimics and subjected to a Mito Stress test on the Seahorse extracellular flux platform, with miR-127-3p, miR-615-3p, and miR-99b-5p included as comparators given their inability to alter circadian period length (Fig. 3F and Fig. 4D, E). All three circadian-active miRNAs reshaped the oxygen consumption rate (OCR) trace, but in clearly distinct directions: miR-27b-3p and miR-191-5p markedly elevated FCCP-induced maximal respiration relative to control, indicating an expansion of spare respiratory capacity (Fig. 4D, top panel), whereas miR-183-5p suppressed maximal respiration below control levels (Fig. 4E, left panel). When resting bioenergetic state was projected onto the OCR–ECAR plane, miR-27b-3p and miR-191-5p shifted myotubes toward an energetic phenotype (Fig. 4D middle panel), while miR-183-5p shifted myotubes toward a glycolytic state (Fig. 4E middle panel). The metabolic-capacity projection, obtained after sequential inhibition of the electron transport chain, reinforced this divergence: miR-27b-3p and miR-191-5p conferred high respiratory capacity (Fig. 4D, lower panel), whereas miR-183-5p confined myotubes to a high-glycolytic capacity (Fig. 4E, right panel). By contrast, the three comparator miRNAs miR-127-3p, miR-615-3p, and miR-99b-5p produced intermediate, largely overlapping bioenergetic profiles relative to NTC, underscoring that the metabolic reprogramming is a selective feature of the circadian-active miRNA subset rather than a generic consequence of transfection (Fig. 4D, E). Together, these data establish that individual PDAC sEV-derived miRNAs are sufficient to induce myotube atrophy, and that the three miRNAs with the greatest impact on the circadian clock additionally remodel mitochondrial respiration and substrate utilization along distinct, non-redundant trajectories.

### Patient-Circulating miRNAs Overlap with the Stable Secretome of Pancreatic Tumor-Derived sEVs

Having established that individual sEV-derived miRNAs reprogram both the molecular clock and the bioenergetic state of muscle cells, we next asked whether the miRNAs identified as active *in vitro* are also present in the circulation of pancreatic cancer patients, and whether they correspond to the miRNAs most consistently secreted by PDAC-derived sEVs over time. Small RNA sequencing was performed on sEVs isolated from the serum of six pancreatic cancer patients (n=4 pancreatic ductal adenocarcinoma, n=1 neuroendocrine pancreatic tumor, n=1 lymphohistiocytoid malignant mesothelioma of the pancreas; Fig. 5A). A total of 428 miRNAs were detected in patient serum sEVs. Unsupervised hierarchical clustering of patient miRNA expression profiles resolved three clusters with distinct signatures (Cluster 1: P1, P3, P5; Cluster 2: P4; Cluster 3: P2, P6; Fig. 5A); the neuroendocrine case (P5) co-clustered with two PDAC samples in Cluster 1, indicating that the cluster structure is driven by the sEV miRNA cargo of pancreatic tumors rather than by histology. Within-cluster Jaccard indices (0.588–0.644) confirmed high intra-cluster similarity (Fig. 5A), although the limited sample size precludes definitive clinical stratification at this stage.

To evaluate the overlap between patient-circulating and tumor-secreted miRNAs, the 428 patient serum miRNAs were intersected with the PANC-1 sEV miRNA dataset. A total of 359 miRNAs were shared between patient serum and tumor-derived sEVs (Fig. 5A, Venn). Stratification by cluster further demonstrated 84.6%, 84.2%, and 83.9% concordance between cluster-specific patient miRNAs and the PANC-1 sEV secretome for Clusters 1, 2, and 3, respectively (Fig. 5A), establishing that the overlap is also robust to inter-patient heterogeneity. UpSet stratification by patient confirmed that the majority of shared miRNAs were detected across multiple patients, indicating that the overlap is not driven by any single individual (Fig. 5B). The expression of the 11 functionally validated candidates across all six patients is displayed in the heatmap below the UpSet plot (Fig. 5B), confirming their consistent representation in patient serum. Ranking of the 20 most abundant serum sEV miRNAs within each patient cluster (Suppl. Fig. 3) further showed that the dominant species, hsa-miR-486-5p, hsa-miR-423-5p, hsa-miR-10b-5p, hsa-miR-22-3p, hsa-miR-10a-5p, hsa-miR-122-5p, hsa-miR-191-5p, hsa-miR-26a-5p, hsa-miR-125a-5p, and the let-7 family, were largely shared across Clusters 1, 2, and 3, with only a small number of cluster-defining species (for example, hsa-miR-143-3p in Clusters 2 and 3, and hsa-miR-451a in Cluster 3). The convergence of the high-abundance fraction across otherwise transcriptionally distinct patient clusters indicates that the dominant serum sEV miRNA signature is a stable feature of the disease, and that the cluster structure resolved by hierarchical clustering arises predominantly from lower-abundance species rather than from a wholesale rearrangement of the cargo.

To extend this observation beyond the cell line and the small serum cohort, we interrogated the pancreatic cancer miRNA-Seq dataset from the GDC data portal (n=495 tumor samples). Because the GDC dataset reports expression at the precursor (hairpin) level, each mature miRNA was mapped to its corresponding precursor(s), and a miRNA was scored as ‘highly expressed’ in a given sample when any of its precursors ranked within the top 50 most abundant miRNAs (Fig. 5C). Ranking within each tumor revealed that hsa-let-7f-5p, hsa-miR-10a-5p, and hsa-miR-27b-3p were among the 50 most abundant miRNAs in more than 90% of samples (99.8%, 94.7%, and 92.3%, respectively; Fig. 5C), identifying them as broadly dominant across the pancreatic cancer landscape. hsa-miR-92a-3p, hsa-miR-26a-5p, hsa-miR-191-5p, and hsa-miR-99b-5p were highly expressed in the majority of tumors (57–76%), whereas hsa-miR-183-5p and hsa-miR-127-3p exhibited enrichment only in a subset of samples, and hsa-miR-30c-5p and hsa-miR-615-3p were rarely or never among the most abundant species (Fig. 5C). The convergence between the miRNAs most consistently secreted by PANC-1 sEVs, those most consistently detected in patient serum, and those most highly ranked across 495 pancreatic tumors indicates that the sEV miRNA signature defined *in vitro* is not an idiosyncrasy of the cell line but a feature of the broader disease.

Across these orthogonal analyses, hsa-miR-27b-3p emerges as the single miRNA satisfying every criterion that the experimental design tests: it is among the three most broadly expressed miRNAs across 495 pancreatic tumors (Fig. 5C); it is consistently detected in patient serum sEVs (Fig. 5B); it is rhythmically expressed in PANC-1 sEVs (cluster 2 in Fig. 3C, D and Supp. Fig. 2); and, in functional assays, it is sufficient to shorten the circadian period and produce a phase shift in the BMAL1 reporter (Fig. 3F, G), to induce myotube atrophy of a magnitude comparable to dexamethasone (Fig. 4C), and to drive an energetic, high-OCR bioenergetic phenotype in muscle cells (Fig. 4D). miR-27b-3p therefore represents a node at which circadian regulation, tumor-derived sEV biology, and cachexia-relevant muscle reprogramming intersect, and a logical priority for mechanistic dissection in vivo.

## Discussion

Pancreatic ductal adenocarcinoma (PDAC) is distinguished among solid tumors by the systemic remodeling that accompanies it, encompassing skeletal muscle wasting, adipose tissue depletion, hepatic metabolic rewiring, immune dysfunction, and disordered glucose homeostasis (Argiles, Stemmler, Lopez-Soriano, & Busquets, 2018). Circadian clock dysregulation is increasingly recognized as an important systems-level feature of PDAC progression that intersects with tumor metabolism, inflammatory signaling, stromal remodeling, and therapeutic resistance (Schwartz et al., 2023; Zheng et al., 2025).

The data presented here position the small extracellular vesicle (sEV) miRNA secretome of the tumor cell as a candidate molecular link. Indeed, EVs have emerged as the principal vehicles by which pancreatic tumors transmit instructive signals to anatomically distant tissues, reprogramming the metabolic and transcriptional state of recipient cells and establishing microenvironments permissive to disease progression (Costa-Silva et al., 2015; G. Wang et al., 2023). PDAC-derived exosomes prime the hepatic pre-metastatic niche through Kupffer-cell activation and stellate-cell fibrotic remodeling, and tumor EVs more broadly drive hepatic metabolic rewiring, including fatty liver formation and suppression of cytochrome P450–dependent drug metabolism, well before the appearance of overt extrahepatic disease. Building on this paradigm, our findings define a distinct, complementary axis of PDAC sEV biology. Whereas the established framework centers on metastasis-supportive niches in the liver and lung, the data presented here identify the host circadian clock and skeletal muscle as previously unrecognized targets of the PDAC sEV cargo: PDAC-derived sEVs dismantle the molecular clock in fibroblasts, osteosarcoma reporter cells, and skeletal myotubes, and induce myotube atrophy in vitro. Critically, the cargo carries time-of-day information, a temporal dimension absent from prior EV-niche models and one that reframes the tumor sEV not as a static delivery vehicle but as a temporally encoded signal capable of imposing pathological timing on recipient tissues. Whether these cell-autonomous effects scale to clinical cachexia is a hypothesis this work motivates rather than establishes.

The clock-disruption phenotype places PDAC alongside the small but growing set of tumors shown to remodel host clocks at a distance. Liver metastases of colorectal origin produce phase shifts in liver and kidney (Huisman et al., 2015), and lung adenocarcinoma rewires hepatic circadian transcription through systemic, tumor-derived signals (Masri et al., 2016). We extend the paradigm to skeletal muscle, a peripheral oscillator whose autonomous BMAL1-driven program orchestrates daily rhythms of mitochondrial function, substrate metabolism, and contractile-protein turnover (Andrews et al., 2010; Hodge et al., 2015; R. A. Martin, Viggars, & Esser, 2023). Particularly informative is the concurrent demonstration by Ducharme et al. that pancreatic-tumor-bearing mice rewire the skeletal-muscle circadian transcriptome through induction of FOXP1, a direct repressor of BMAL1 (Ducharme et al., 2025). The cell-intrinsic FOXP1 program and the cell-extrinsic, sEV-encoded program documented might engage the muscle BMAL1 axis at different molecular layers, suggesting that protection of the muscle clock will require interventions addressing both arms.

A central conceptual point of the present work is that the sEV miRNA cargo resolves into two functionally distinct pools. Sorting of miRNAs into EVs occurs through both selective and non-selective mechanisms operating on biochemically distinct vesicle subpopulations (Temoche-Diaz et al., 2019), with a broad family of RNA-binding proteins and primary-sequence motifs (EXOmotifs) governing the selective arm (Garcia-Martin et al., 2022). The temporal dimension of this regulation has so far been demonstrated most directly at the protein level. Yeung et al. showed in synchronized tendon fibroblasts that individual sEV protein abundances oscillate over 24 h, with subpopulations enriched in RNA-binding proteins temporally separable from those enriched in matrix and cytoskeletal proteins (Yeung et al., 2022). The 67 MetaCycle-rhythmic miRNAs and the temporally defined clusters identified in PANC-1 sEVs extend that protein-level finding to the miRNA fraction in a tumor context. A counterexample from a non-pathological context, no circadian variation in milk-EV miRNA cargo of dairy cows (Saenz-de-Juano, Silvestrelli, & Ulbrich, 2023), tempers any claim of universality and indicates that rhythmic cargo loading is context-dependent, plausibly amplified in pathological cell states.

The bioluminescence and RT-qPCR readouts of recipient-cell clock function diverged in the directionality of the period change. The two assays interrogate distinct regulatory layers, transcriptional initiation versus steady-state transcript pool, and the opposite directionality therefore identifies a post-transcriptional contribution to clock remodeling. miRNAs are the parsimonious candidate, and the now-extensive literature on miRNA control of canonical clock components, including miR-192/194 on Period (Nagel, Clijsters, & Agami, 2009), miR-24/-25-3p/-30a on PER2 translation and/or PER2 stability (Park et al., 2020; Yoo et al., 2017), and miR-142-3p on Bmal1 (Shende, Neuendorff, & Earnest, 2013), provides a mechanistic precedent.

Among the eleven miRNAs tested, miR-27b-3p produces the largest combined effect on circadian period and phase, and drives myotube atrophy of a magnitude approaching the dexamethasone control. The mechanistic embedding of miR-27b-3p in clock and muscle biology has independent support. miR-27b-3p oscillates in metabolic tissues, notably mouse liver, and directly represses BMAL1 by binding its 3⍰UTR, with functional consequences for BMAL1 protein rhythms and hepatic metabolism (W. Zhang et al., 2016). In skeletal muscle, miR-27b targets Pax3 to gate myogenic specification (Crist et al., 2009) and myostatin to regulate proliferation, differentiation balance (Ling et al., 2018). The bioenergetic phenotype we document, elevated maximal respiration and an energetic OCR–ECAR signature, is in apparent tension with the report that miR-27b targets Foxj3 in C2C12 myocytes and reduces mitochondrial biogenesis through the Foxj3–Mef2c–PGC1α axis (Shen et al., 2016), indicating that target prioritization is context-dependent in mature myotubes.

The reciprocal phenotype is mechanistically informative. Transfection of myotubes with miR-183-5p mimic suppressed maximal mitochondrial respiration and shifted substrate utilization toward glycolysis. Wang et al. showed that loss of miR-183/miR-96 in skeletal muscle relieves FoxO1- and PDK4-mediated repression of glucose oxidation and enhances oxidative capacity (H. Wang et al., 2021); the reciprocal prediction, that gain of miR-183-5p suppresses oxidative phosphorylation, is precisely what our gain-of-function data show. All eleven candidate miRNAs nonetheless induce significant myotube atrophy at 48 h, and GO enrichment of their validated targets distributes across ubiquitin–proteasome, autophagy, mitochondrial-organization, and skeletal-muscle-differentiation programs. The cargo operates as a multi-pronged program rather than through a single dominant effector, a redundancy that may help explain why blockade of any single circulating cachexia mediator has not, to date, proved clinically tractable (Paval et al., 2022; Roeland et al., 2020).

Two orthogonal datasets support the translational relevance of these findings. First, 84% of the miRNAs detected in serum sEVs from six pancreatic-cancer patients were also present in the PANC-1 sEV secretome, indicating that the miRNAs identified in vitro circulate in patient blood. Second, let-7f-5p, miR-10a-5p, and miR-27b-3p ranked among the top 50 most abundant miRNAs in over 90% of 495 pancreatic tumors profiled by miRNA-Seq, establishing that these effectors are not idiosyncratic to PANC-1 but broadly expressed across the disease. The relevance of EV-miRNA signaling may extend beyond PDAC: exosomal miR-191-5p, also rhythmically packaged in our dataset, has independently been identified as a functional biomarker in esophageal squamous cell carcinoma (H. Wang et al., 2025). Capturing the rhythmic effectors this work identifies will, however, require time-of-day-resolved sampling, a methodological prerequisite that single-time-point biomarker studies systematically obscure.

An integrated mechanistic model emerges from these observations (Fig. 5D). Within the PDAC tumor cell, a dysregulated clock (BMAL1↓, altered rhythmic output) is coupled to selective miRNA sorting at the site of EV biogenesis: miR-27b-3p is rhythmically packaged into the secreted vesicle pool, whereas the remaining functionally validated miRNAs are loaded constitutively, defining two cargo pools that are temporally distinct at the source yet jointly delivered to the recipient cell. Secreted sEVs are stable in size but variable in number across the day, and the resulting cargo is detectable in patient serum, with 84– 85% concordance against the PANC-1 secretome across the three patient clusters. In the skeletal muscle cell, the integrated cargo dismantles the muscle clock at the levels of period, phase, and amplitude, and concurrently drives bioenergetic remodeling along four non-redundant trajectories: an energetic phenotype (miR-27b-3p and others; ↑OCR, aerobic/oxidative, ↑spare respiratory capacity), a high-metabolic phenotype (miR-191-5p and others; ↑OCR, ↑ECAR, high respiratory capacity), a quiescent phenotype (other miRNAs, ↓OCR, ↓ECAR, low metabolic capacity), and a glycolytic phenotype (miR-183-5p and others; ↓OCR, ↑ECAR, glycolytic drift). The co-occurrence of clock disruption and bioenergetic remodeling within the same recipient cell type, together with the predicted (target-enrichment-based) engagement of proteostatic networks, is consistent with the myotube-atrophy phenotype documented here *in vitro*, although the present data do not establish a direct causal hierarchy among these processes nor demonstrate that any of them is individually necessary or sufficient for atrophy in this system. Whether these molecularly distinct perturbations integrate, *in vivo*, into the systemic phenotype of pancreatic cancer cachexia, and, if so, whether they do so additively, synergistically, or through a dominant arm that the current *in vitro* readouts do not separate, remains an experimentally tractable hypothesis rather than an established conclusion. Testing this hypothesis will require tumor-bearing animal models in which sEV release, miRNA cargo composition, and time-of-day delivery can be perturbed independently, with muscle clock function, mitochondrial respiration, proteostatic flux, and overt mass loss measured as separable endpoints. Within that framework, the model proposed here positions the temporally encoded, multi-channel sEV signal as a candidate organizing principle for cachexia rather than as a single-effector mechanism, and provides a molecularly defined entry point, miR-27b-3p in particular, against which loss-of-function and chronopharmacological interventions can be benchmarked.

Beyond the *in vivo* program outlined above, two further dimensions of the present work define a path for follow-up investigation. The small-RNA sequencing performed on PANC-1 provides a high-resolution temporal map of a representative PDAC line, and its extension to additional cell lines and to the histologically heterogeneous patient cohort, particularly through longitudinal sampling, will determine the breadth of the rhythmic program across the disease spectrum. In parallel, time-of-day-resolved profiling of patient serum EVs, beyond the single-time-point sampling that the present cohort necessarily reflects, will be required to establish whether the rhythmic miRNA architecture defined *in vitro* is preserved at the systemic level in cancer-bearing patients, a question that the cell-line data raise but cannot, on their own, resolve.

Three implications follow from these findings. The most immediate is methodological: EV-based biomarker programmes in PDAC will benefit from time-of-day-resolved sampling, since rhythmically packaged miRNAs are systematically obscured by single-time-point analysis and represent a class of effectors that current biomarker pipelines are not designed to capture. The translational corollary (Fig. 5D) can be made that if the secretory acrophase of miR-27b-3p, or of other rhythmic effectors, falls within a circumscribed daily window, antimiR delivery or sEV-uptake blockade could be timed to that window, in principle reducing systemic exposure while preserving therapeutic efficacy, a chronopharmaceutical logic that single-time-point pharmacology cannot exploit. Most broadly, the rhythmic structure documented here in PDAC raises a biological question whose scope extends well beyond pancreatic cancer, whether comparable temporal organization characterizes the EV cargo of other malignancies, and whether it engages the same recipient-tissue programs. The methodology developed in this work is positioned to address that question, and its answer will determine whether circadian regulation of tumor-derived EV signaling is a feature specific to PDAC or a general principle of cancer-host communication.

## Author Contributions

**Jonathan Church**: conceptualization, methodology, investigation, formal analysis. **Ignacio Aiello:** methodology, investigation, formal analysis, visualization, validation, writing – review and editing. **Alessandro Ceci**: investigation, formal analysis. **Goeun Jang:** investigation. **Carla V. Finkielstein:** conceptualization, supervision, funding acquisition, project administration, resources, writing – review and editing, visualization.

## Acknowledgements

The authors acknowledge the members of the Finkielstein laboratory for their valuable discussions, feedback, and technical support throughout this study. The authors also thank the Englander Institute for Precision Medicine at Weill Cornell Medicine for assistance with patient sample collection and clinical coordination.

## Geolocation Information

This work was completed at the Fralin Biomedical Research Institute at VTC, 2 Riverside Cr., Roanoke, Virginia 24016, US.

## Data Availability Statement

The small-RNA sequencing data generated in this study and all other data supporting the findings of this study, including bioluminescence time-courses, RT-qPCR measurements, myotube-diameter quantifications, extracellular-vesicle characterization data, and Seahorse extracellular-flux traces, are available from the corresponding author upon reasonable request. Patient-derived sequencing data are subject to controlled access in accordance with the terms of the Virginia Tech and Weill Cornell institutional review board approval and patient informed consent.

## SUPPLEMENTARY MATERIAL

**Supplementary Figure 1. Clock-Gene Expression In Circadian-Synchronized PANC-1 cells.** RT-qPCR analysis of BMAL1, CRY2, DBP, NR1D1, and PER2 expression in synchronized PANC-1 cells harvested at indicated time-points following dexamethasone synchronization. MetaCycle analysis (Meta2d) showed no statistically significant rhythmicity for any of the five core clock transcripts (table inset).

**Supplementary Figure 2. Rhythmically Expressed miRNAs in PANC-1 sEVs.** (**A**) Heatmap of the 67 miRNAs detected as significantly rhythmic by MetaCycle (p<0.05) in sEVs collected every 4 h for 36 h, ordered into four phase windows: Early (0–8 h, blue), Mid-day (8–16 h, yellow), Late (16–24 h, orange), and Night (>24 h, purple). Significance categories (p<0.01; 0.01–0.025; 0.025–0.05) are indicated by the color band on the left. (**B**) Rayleigh plot of the 67 rhythmic miRNAs. The angular position of each point indicates the acrophase (peak expression time) calculated as 2π × phase/period, mapped onto a 24-hour clock (0 h at top). The radial distance represents relative oscillation amplitude, normalized to the range of all detected amplitudes (low at center, high at periphery). Point color reflects the estimated oscillation period, from short (~18.8 h, purple) to long (~27.9 h, yellow). The black arrow represents the Rayleigh mean vector; its length (R) indicates the degree of phase clustering across all oscillating miRNAs (R=0, uniform distribution; R=1, perfect clustering).

**Supplementary Figure 3. Top 20 miRNAs Per Patient Cluster Ranked by Mean Expression in Serum sEVs.** Bar plots of the 20 most highly expressed miRNAs in each patient cluster, ranked by mean log2 expression: Cluster 1 (red, top), Cluster 2 (blue, middle), and Cluster 3 (green, bottom).

**Supplementary Table 1. MetaCycle Analysis Of Clock Genes Rhythmicity in NIH3T3 and C2C12 Cells.** Circadian rhythm parameters (p-value, period, phase, mesor, and amplitude) for *Bmal1, Per2*, and *Cry2* expression in NIH3T3 and C2C12 cells under control conditions and following treatment with 100% PANC-1 cell-conditioned media (CM). This analysis quantifies the temporal oscillatory patterns shown in Figure 1G-I and J-L.

