## Supplementary figures and images for "Pancreatic cancer extracellular vesicles carry a time-of-day-regulated miRNA cargo that disrupts the skeletal muscle clock and bioenergetics"

### Supplementary Figure 1

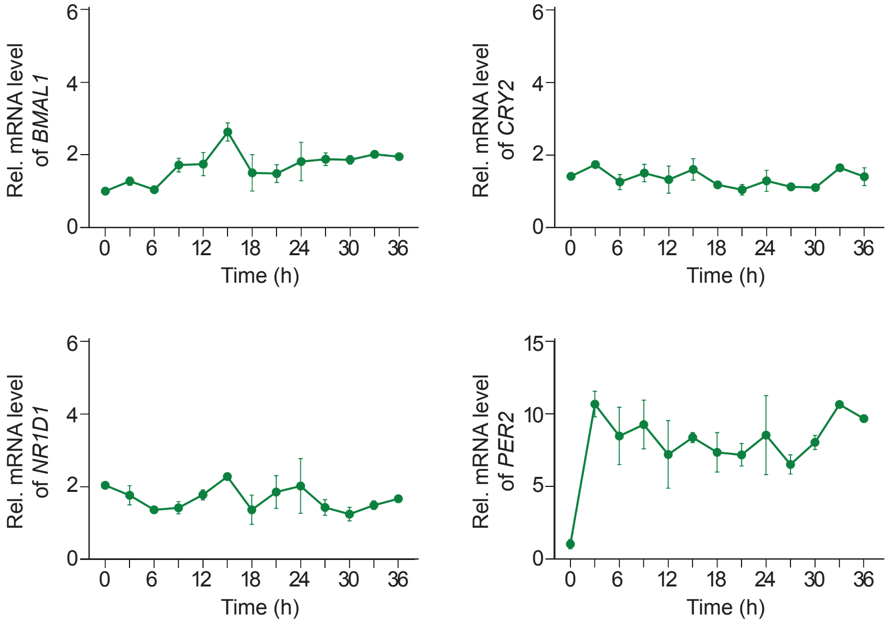


| | Meta2d | | | | |
| --- | --- | --- | --- | --- | --- |
| PANC1 cells | Gene | p value | Period | Phase | Amplitude |
| | *BMAL1* | 0.170 | 22.739 | 13.727 | 0.179 |
| | *PER2* | 0.754 | 25.196 | 9.229 | 0.204 |
| | *CRY2* | 0.759 | 27.489 | 6.414 | 0.105 |
| | *NR1D1* | 0.203 | 22.987 | 18.017 | 0.151 |
| | *DBP* | 0.702 | 26.937 | 22.381 | 0.124 |

### Supplementary Figure 2

Sup_Fig_2_Church_Aiello_et al.

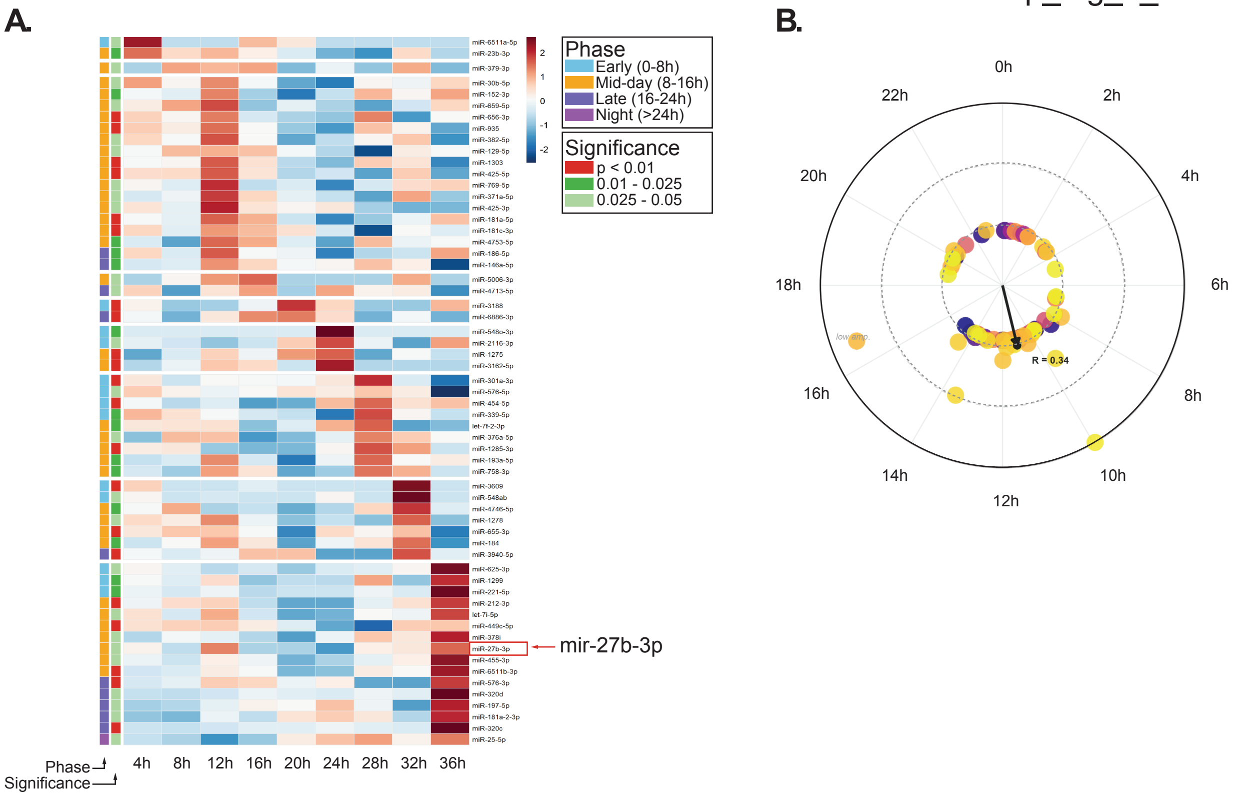

### Supplementary Figure 3

Sup_Fig_3_Church_Aiello_et al.

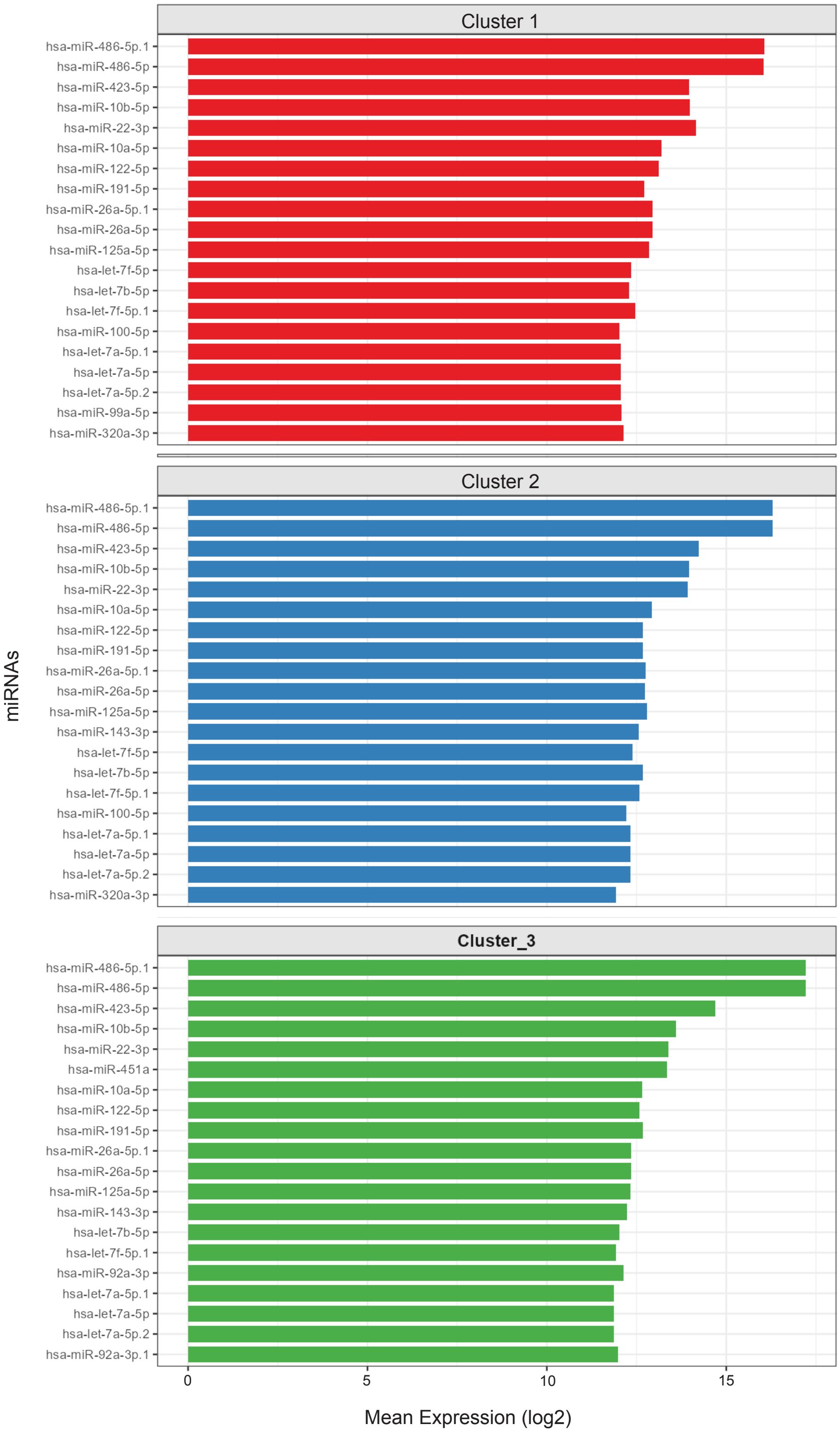
