## Supplementary Table 1 for "Pancreatic cancer extracellular vesicles carry a time-of-day-regulated miRNA cargo that disrupts the skeletal muscle clock and bioenergetics"

**Supplementary Table 1. MetaCycle analysis of clock genes rhythmicity in NIH3T3 and C2C12 cells.**

|  | **Gene** | **p value** | **Period** | **Phase** | **Amplitude** |
| --- | --- | --- | --- | --- | --- |
| **NIH3T3 cells** | *Bmal1* control | 9.86E-07 | 25.93 | 8.22 | 0.96 |
|  | *Bmal1* CM | 1.09E-05 | 27.66 | 10.49 | 1.26 |
|  | *Per2* Control | 6.56E-05 | 24.1 | 0.81 | 0.71 |
|  | *Per2* CM | 3.87E-07 | 25.11 | 24.26 | 0.57 |
|  | *Cry2* control | 0.2891 | 27.57 | 24.15 | 0.53 |
|  | *Cry2* CM | 0.0359 | 27.087 | 21.25 | 0.41 |
| **C2C12 cells** | *Bmal1* control | 0.0135 | 19.99 | 9.44 | 0.41 |
|  | *Bmal1* CM | 8.60E-01 | 25.33 | 4.15 | 0.14 |
|  | *Per2* Control | 3.81E-03 | 25.35 | 22.06 | 0.37 |
|  | *Per2* CM | 0.0163 | 20.43 | 0.76 | 0.44 |
|  | *Cry2* control | 0.0387 | 23.25 | 22.62 | 0.19 |
|  | *Cry2* CM | 0.2554 | 23.1 | 22.46 | 0.29 |
